# Generative design of *de novo* AAV vectors with unique serological profiles

**DOI:** 10.64898/2026.11.06.764881

**Authors:** Giulio Chiesa, Suyue Lyu, Maria D. Nguyen, Jared Benjamin, Michael Garton

## Abstract

More than 5000 monogenic diseases are potentially curable by replacing a single defective gene with one that functions normally. Currently, the only clinically approved vector able to effectively deliver a therapeutic gene to a target tissue is Adeno-Associated Viruses (AAVs). However, despite decades of work, only a handful of gene replacement therapies have been approved, and many patients remain ineligible for even those. A major barrier to broad adoption of AAV therapies is pre-existing immunity to AAVs, mediated by neutralizing antibodies (nAbs). It is potentially possible to change exposed capsid surfaces in AAV capsids to avoid nAb recognition. However, traditional techniques such as rational design and directed evolution only allow modest changes. AAV function becomes compromised long before full nAb avoidance is achieved. Recent advances in protein design by generative modelling provide a new way to overcome this problem by learning AAV design constraints from natural sequences. We built a variational autoencoder that could learn these sequence constraints and generate functional AAV capsid proteins *de novo*. More than 255 designs were synthesized and tested *in vitro* in two design sets, with one set dedicated to exploring uncertain regions of the sequence space. Between the two campaigns, seven designs transduced human cells with >50% efficiency, of which four avoided nAb neutralization (NT50 < 1:50) in all human sera that were reactive to AAV. In one of these designs, sequence identity to the nearest natural AAV sequence was 18% divergent. Given the radical sequence diversity and its unique serological profile, this new capsid design would represent the first ever AAV serotype designed *de novo*.

## Introduction

With seven approved therapies currently being administered in clinic and hundreds more in clinical trial^1^, gene replacement therapy by means of Adeno-Associated Viral (AAV) vectors is one of the most promising platforms for widespread treatment of hereditary disease such as Retinal Dystrophy, Spinal Muscular Atrophy, hemophilia A and B, and Duchenne Muscular Dystrophy^1^. AAV are attractive because they have relatively low immunogenicity and pathogenicity compared with other vectors such as adenovirus^2^. They can also efficiently target and transduce multiple clinically important tissue types and sustain high levels of persistent payload expression over long time scales and lower genome toxicity^3^.

Despite these advantages, one of the major barriers to the broader development of AAV therapies is the pre-existing humoral response to AAVs, which results from environmental exposure and raises safety concerns while reducing efficacy^4,5^. This is amply illustrated by Haemophilia A patients, where almost 60% have pre-existing neutralizing antibodies to AAV2 capsid epitopes^6^. As a result, large segments of patient populations can be excluded from eligibility^7^.

Several strategies have been developed to address this, from rational design ^8–11^ to ancestral sequence reconstruction and directed evolution under immune pressure^12^. These approaches, however, remain largely confined to near-natural sequence space, unlikely to produce viral-surface-wide mutations needed for broadly avoiding pre-existing nabs across different patients^8–11^.

Recently, deep generative models designed functional proteins across many families with higher diversity. These models could potentially produce functional capsid proteins that are more distant from natural serotypes. Previous applications in AAV have mostly focused on one region of 28 amino acids, with abundant functional annotations for various mutations, and viable capsids with 12–29 mutations in this region successfully generated^13–15^. However, most epitopes recognized by mAbs lay outside of the region addressed in these studies. To date, only one method has been experimentally validated to produce viable VP3 with a large number of mutations (up to 30% of residues) across different sequence regions. In this recent work, the AAV2 VP3 capsid protein was redesigned using ProteinMPNN. However, changes at the viral surface positions that typically mediate nAb recognition were explicitly avoided, and it is not clear that this approach could be applied to develop AAV variants able to resist nAb neutralization^16^. Therefore, a new method that introduces mutations across the capsid surface while retaining assembly and transduction is required for evading pre-existing nAbs.

To fill this gap, we developed a new variational autoencoder model, and experimentally evaluated generated sequences. We modelled only the VP3 protein, because it forms most of the external capsid surface and carries dominant nAb epitopes^17^. We previously developed a variational autoencoder (ProteinVAE) computationally validated on adenovirus hexon sequences, a problem with comparable challenges, including long sequences and limited samples^18^. Here we adapt ProteinVAE for the design of AAV capsids that avoid pre-existing nAbs against natural AAVs. With an additional tailored contrastive loss term, ProteinVAEc was specifically designed to navigate the imbalanced and low diversity natural AAV sequence space. A total of 255 variants were generated *de novo*, ranging from 5 to 99 mutations relative to the closest natural sequence and distributed across VP3. Seven designs successfully transduced multiple human cell types (> 50%). Nine designs avoided nAb mediated neutralization (NT50 < 1:50) in by human blood serum that neutralizes multiple natural AAV serotypes. Four nAb evading variants had > 50% transduction efficiency, and seven had >40%. One design, that transduced HEK293T cells with 64% efficiency, had 97 mutations compared to the closest natural sequence and was capable of broad evasion across all sera tested.

Altogether, this represents the first example of experimentally validate *de novo* AAV design that yields capsids with a different serological profile than their closest neighbor, thus representing the first ever *de novo* new serotype.

## Results

To explore AAV capsid sequence space while preserving functionally important constraints, we curated 355 full-length VP3 sequences from AAVs and related dependoparvoviruses from Uniprot database. The dataset is strongly imbalanced with 51% of sequences sharing at least 75% identity with AAV2. To improve representation learning on this challenging dataset, we added a tailored contrastive term in the loss. We refer to the resulting model as ProteinVAEc. All other architectural and training settings were identical to ProteinVAE.

### Contrastive learning improves sequence quality

To assess learning of natural VP3 patterns, ProteinVAEc, was first evaluated against both ProteinVAE and fine-tuned ProGen2 models using sequence and structure metrics *in silico*. Sequences were unconditionally sampled and top likelihood sequences were used for comparison (see Methods). Generated sequences from ProteinVAEc showed the lowest column-wise Jensen–Shannon divergence (JSD) from the natural VP3 alignment, indicating better recovery of evolutionary constraints (**Figure 1a**). Predicted secondary structure content also showed that helix and strand fractions in ProteinVAEc sequences matched natural VP3 proteins more closely (**Figure 1b**). In addition, ProteinVAE sequences populated only the low-helix region, whereas ProteinVAEc additionally recovered a high-helix cluster present in natural VP3, indicating that the contrastive learning mechanism preserved a mode of natural structural variation that the baseline model failed to capture.

Generated VP3 can be used to form a mosaic capsid with natural VP1 and VP2. However, AAV capsid proteins cannot be freely combined across serotypes because structural divergence can cause incompatibilities; for example, AAV4 and AAV5 VP3 proteins are not interchangeable with AAV2 VP3^19–21^. Therefore, a reasonable search space for synthetic VP3, with reduced risk of structural clashes, is between pairs of known compatibility. The variational autoencoder architecture of ProteinVAEc offers latent interpolation as a convenient way to search this space. To demonstrate this use case, sequences were interpolated between AAV1 and AAV2 with both ProteinVAE and ProteinVAEc. ProteinVAE produced irregular, non-monotonic changes in Hamming distance, whereas ProteinVAEc produced a smooth and monotonic transition toward both AAV1 and AAV2, consistent with a smoother latent space (**Figure 1c,d**). This smoothness also allows predictable control over how different the generated sequence is from each endpoint. Next, fine-tuned ProGen2 likelihood was used as an independent metric for sequence plausibility, with ProteinVAEc interpolation achieving higher likelihood than ProteinVAE (**Figure 1e**). Overall, these findings show that the tailored contrastive learning mechanism improved representation quality and produced sequences with structure and patterns closer to those of natural sequences, and therefore a higher likelihood of retaining function. With promising *in silico* evaluation results, we selected sequences for experimental validation.

**Figure 1.**
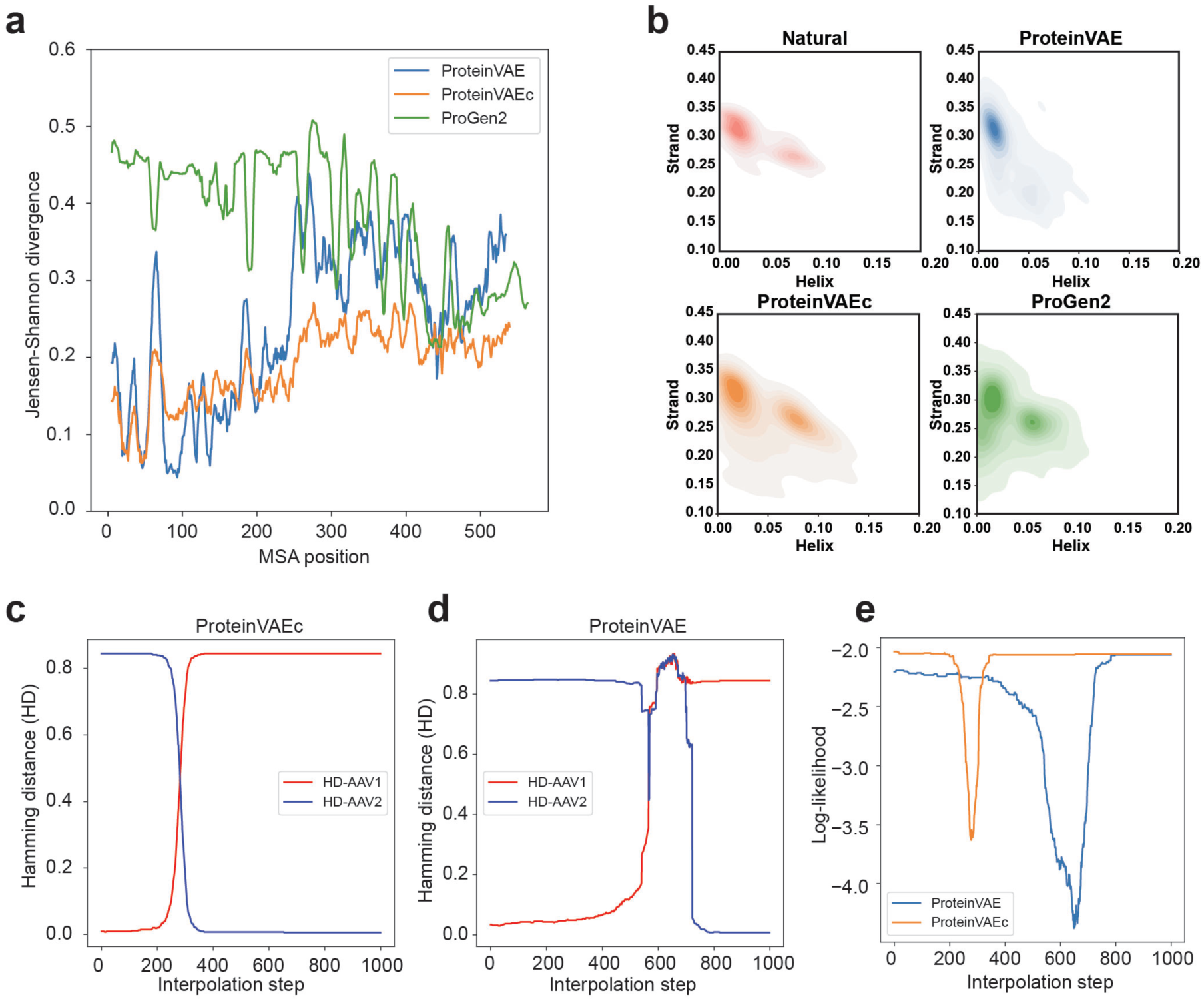
ProteinVAEc shows improved recovery of natural sequence statistics compared with ProteinVAE, while both versions outperform fine-tuned ProGen2. **(a)** MSA column-wise JSD between natural and generated AAV VP3 sequences. Jensen– Shannon divergence across MSA columns between natural sequences and sequences generated by different models. Lower JSD indicates closer agreement with natural amino acid distributions. ProteinVAEc demonstrates improved recovery of natural sequence statistics compared with ProteinVAE, while both versions outperformed fine-tuned ProGen2. **(b)** Helix and strand composition of natural and generated sequences; ProteinVAEc produces secondary-structure distributions that more closely match natural sequences than those generated by ProteinVAE. **(c)** Sequences generated by ProteinVAE exhibit irregular, non-monotonic changes in Hamming distance relative to the two endpoint sequences. **(d)** In contrast, ProteinVAEc generates interpolated sequences with monotonic progression in Hamming distance toward each endpoint, consistent with smoother traversal of latent space. (**e**) ProteinVAEc sequences also achieve higher likelihood under the fine-tuned ProGen2 model than ProteinVAE, indicating improved compatibility with evolutionary and structural constraints.

### Experimental validation identified functional variants with up to 18% sequence divergence from closest natural sequence

To evaluate generated sequences experimentally, two sets of synthetic VP3 proteins were generated with two distinct objectives: (1) low risk exploitation; and (2) high risk exploration (**Figure 2a**). As the VP1/2 unique regions were supplemented from AAV2 in this experimental validation, the local region of AAV2 in latent space was prioritized. A set of 100 variants (Set 1) was generated by ProteinVAEc by sampling a Gaussian distribution centered on AAV2 and interpolating between AAV2 and its known compatible AAVs. Decoded sequences were then filtered by additive fitness score with sequences containing deleterious mutations removed (see Methods). Mutation load in Set 1 was limited to 2-10 mutations from the closest natural sequence (AAV2, **Figure 2b**). Variants contained mutations across the whole VP3 region, including folded regions, loops and hypervariable regions (HVRs).

**Figure 2:**
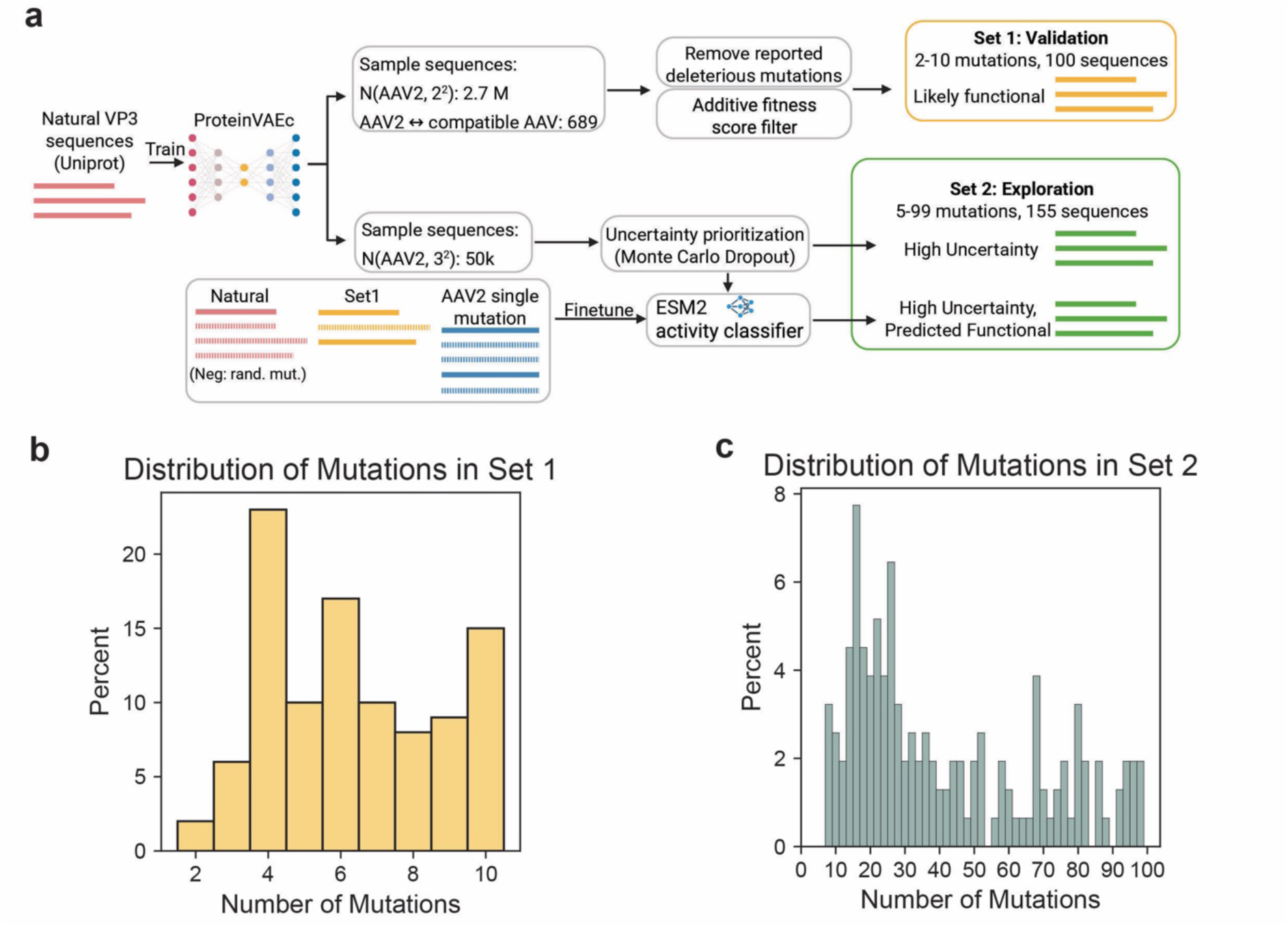
Synthetic VP3 variants carry mutations across every VP3 functional region. (**a**) Schematics of *in silico* screening of generated sequences. ProteinVAEc generates candidate VP3 sequences from the AAV2 latent region. Set 1 (2–10 mutations) is sampled near AAV2 and variants with double mutations at R585 and R588 or mutations in regions involved in AAV2–AAVR interaction were removed. Remaining variants were filtered through an additive fitness criterion from previous single mutation biodistribution data (see Methods). Set 2 (exploration, 5–99 mutations) is sampled further from AAV2 and prioritized first by Monte-Carlo-dropout uncertainty, then filtered with fine-tuned ESM2 activity classifier. **(b, c)** Number of mutations in Set 1 (**b**) and Set 2 (**c**).

A second set comprising 155 variants (Set 2) was generated by sampling a Gaussian with larger standard deviation (see Methods). Uncertainty was quantified with Monte Carlo Dropout method, and uncertain regions of the latent space were prioritized to get more informative samples (see **Methods**). To increase functional fraction, sequences from these regions were further prioritized according to their predicted activity by a fine-tuned ESM2 classifier. The number of mutations in Set 2 VP3 variants ranged from 5 to 99 compared to the closest natural sequence of AAV2 (**Figure 2c**). Variants of both sets were individually cloned and produced in HEK293T LentiX cells. Payload comprised an EGFP gene under a CMV early enhancer / Chicken beta-Actin (CAG) promoter (**Figure S1**) to enable titer measurement by quantitative PCR (qPCR) and transduction efficiency measurement by flow cytometry of EGFP+ cells (**Figure 3a**, gating strategy in **Figure S2**). Batch production was chosen over a library approach to avoid cross-packaging^22^.

To evaluate different aspects of VP3 function, synthetic variants were tested for *in vitro* transduction, capsid assembly, and genome packaging. Selection criteria were applied to promote promising candidates for more costly evaluation steps. First, transduction efficiency was measured in HEK293T with a fixed volume of viruses (*transduction screening step*, **Figure 3b, c**), where number of virus particles can vary across variants. Then, viral titre and packaging capacity were evaluated in parallel for each variant included in the first screening (**Figure 3d, e**). Lastly, promising variants were selected to be produced in higher quantity for transduction evaluation at equal number of Multiplicity of Infection (MOI = 500) in HEK293 cells, to decouple packaging efficiency and transduction efficiency (*transduction validation step*, **Figures 3f** and **S6a, b**).

In the transduction screening step, 27 variants of Set 1 transduced HEK293T cells with an efficiency >30%. Of these, 19 had a transduction efficiency of >50% of cells, and seven were comparable to AAV2 levels (**Figure 3b**, red dots). In Set 2, 155 variants were screened, but only four variants transduced better than the PBS control (one-sided Welch’s t-test on log EGFP+%, BH q < 0.05; **Methods**). Three variants transduced >10% cells (respectively, s2_131, s2_0 and s2_129). Of these three variants, one carried 11 mutations (s2_131), while the other two carried respectively 93 (s2_0) and 97 mutations (s2_129), representing ∼18% of sequence diversity from AAV2.

In the qPCR screening step, a high throughput method for measuring packaging competence was adapted^23^. Incomplete packaging is a critical pitfall in AAV production caused by incomplete load of the viral genome^24,25^ or by contamination of parts of the packaging plasmids or genomic DNA^24^. It can severely reduce transgene expression^25^. Probes targeting the sequence of either EGFP or ITR were adopted in parallel reactions^23^. A standard curve for each titre was generated by including serial dilutions of a commercially obtained AAV2 stock solution in each plate (Representative curve in **Figure S4**). Upon conversion, titers were expressed as the log10 of viral genome per microliter, or vg/µl (**Figure 3d, e**). Variants were classified as *virus producing* when converted ITR titreexceeded the PBS background control by at least 3 Standard Deviations, while they were considered *packaging competent* by applying the same threshold to log10 EGFP titre. In Set 1, of 100 variants, 78 are *virus-producing* (mean log10 ITR titer more than 3 SD above PBS), and 20 of those 78 were considered *packaging competent* (**Figure 3d**). A negative correlation was found both between number of mutation and EGFP titre (Spearman rho=-0.483, p=3.65e-07) and number of mutation and ITR titre (Spearman rho=-0.345, p=4.33e-04), suggesting that higher number of mutations increases the chances of disruption of either the folding, assembly, (low ITR titre) or the packaging machinery of the capsid (low EGFP titre).

In Set 2 (**Figure 3e**), 68 variants out of 155 had ITR titres higher than threshold (*virus producing*), while 23 of *virus producing* variants had EGFP titres higher than threshold (*packaging competent*). A negative correlation was again found between number of mutations and EGFP or ITR titres, albeit to a lower degree (for ITR: Spearman rho=-0.157, p=5.08e-02;); for EGFP: Spearman rho=-0.294, p=2.09e-04). Lastly, the EGFP/ITR ratio was calculated for variants considered *packaging competent* for each set (**Figure S5a, b**), but no clear correlation was detectable between EGFP/ITR ratio and number of mutations. Overall, qPCR data suggest that designed variants fold correctly (hence protecting DNA in the DNAseI reaction), but are often affected by packaging defects, as demonstrated by the lower EGFP titres across both sets.

Among Set 1 variants, those that transduced >50% of HEK293 cells in the first screening (**Figure 2c**) were selected for production scale up. DdPCR confirmed that 19 out of 22 variants had titres ≥4 × 10^5 vg/µL, which was sufficient for testing at 500 MOI. The three lower-titre variants were excluded. A second subset of variants that showed high titres in the qPCR experiment was selected at random for scale up production and included in the experiment (**Figure S6a**). For Set 2, any variant with titre ≥4 × 10^5 vg/µL was selected (77 variants, **Figure 3f** and **S6b**). At MOI of 500, four variants from Set 1 transduced with >50% efficiency (**Figure S6a**), while only one variant of Set 2 (s2_129, 96 mutations) had a ∼64% transduction efficiency. Among the remainder of Set 2 variants, 23 transduced >1% cells, of which five transduced >10%, confirming that Set 2 contains overall less transduction competent variants.

**Figure 3.**
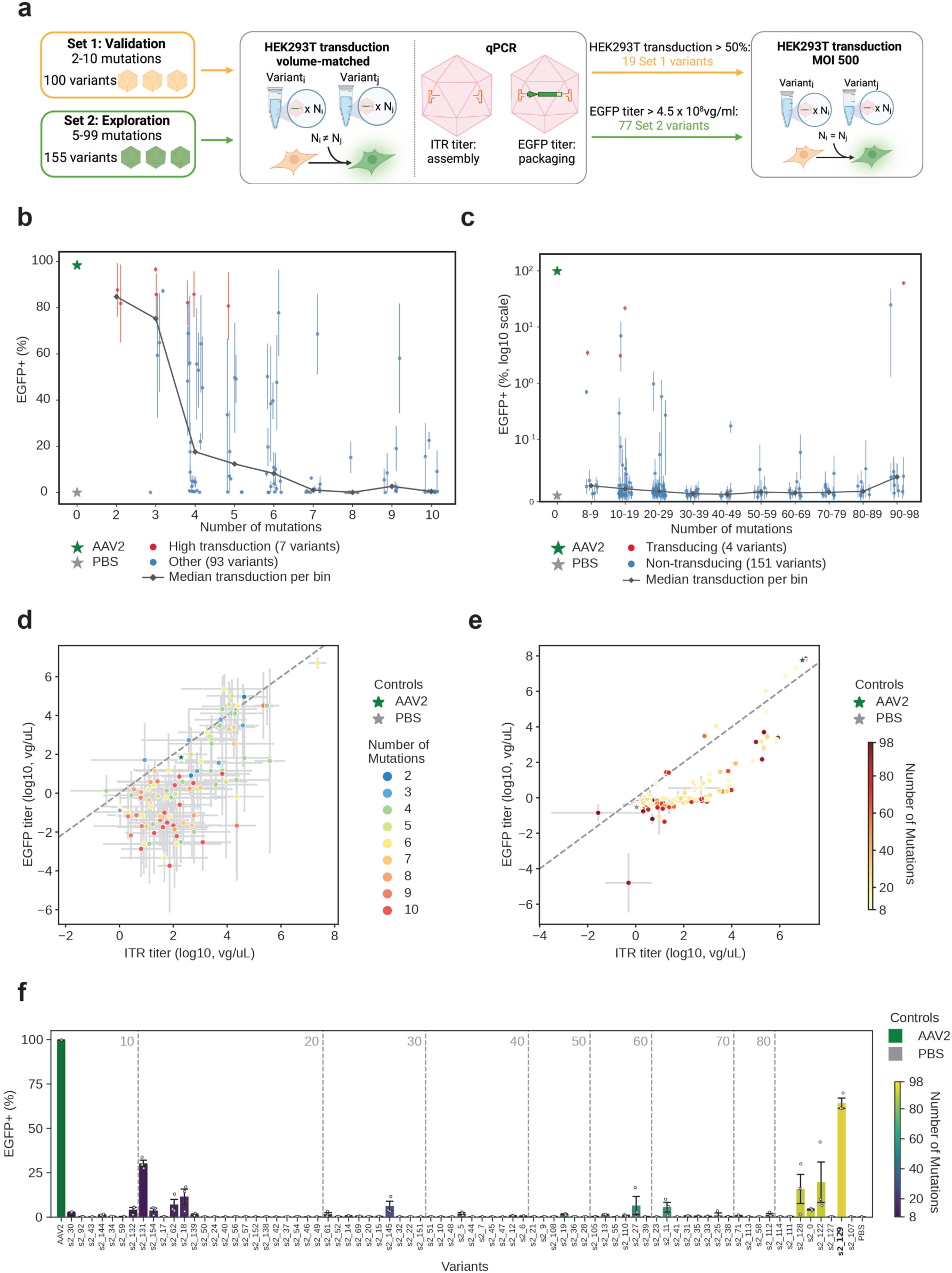
Arrayed screening for transduction and packaging identifies transduction competent variants with up to 96 mutations. (**a**) Schematics of the selection step, from transduction screening to validation. All variants are tested in a transduction screening with matched volumes (**b, c**) and a parallel qPCR screening with EGFP and ITR probes (**d, e**). Selected variants from the transduction and the qPCR screenings are then tested in a new transduction screening at 500 MOI (**f, S6, S7**). **(b)** HEK293T transduction efficiency (volume-matched) vs. mutation count for all 100 Set 1 variants. Y axis shows the mean EGFP+ cell percentage. X axis shows the mutation count against AAV2 (see **Methods**), with horizontal jitter for visibility. Error bars are SEM across three biological replicates. Seven variants (red) were classified as high-transducing with EGFP+ > 80% while remaining not significantly below AAV2 (one-sided Welch’s t-test on log EGFP+%, BH q ≥ 0.05; Methods). Variants not classified as high-transducing are shown in blue. Black line shows median EGFP+% per mutation count. Wild-type AAV2 (dark green star) and PBS (grey star) are used as positive and negative controls, respectively. **(c)** HEK293T transduction efficiency (volume-matched) vs. mutation count for all 155 Set 2 variants. Y axis shows the mean EGFP+ cell percentage on log10 scale. X axis shows the mutation count against AAV2 VP3 (see Methods), with horizontal jitter for visibility, binned every 10 mutations. Error bars are SEM across three biological replicates. Four variants (red) were classified as transducing, defined as EGFP+ significantly higher than PBS (one-sided Welch’s t-test on log-transformed EGFP+%, Benjamini–Hochberg adjustment, q < 0.05; see Methods). Variants not classified as transducing are shown in blue. Black line shows median EGFP+% per mutation count bin (every 10 mutations). Wild-type AAV2 (dark green star) and PBS (grey star) are used as positive and negative controls, respectively. (**d,e**) Viral titres measured by qPCR using ITR (x-axis) and EGFP (y-axis) probes, expressed as log₁₀(vg/µL). Conversion from ΔCq values is described in **Methods**. Wild-type AAV2 (green star) and PBS (grey star) are included as positive and negative controls. Variant color indicates the number of mutations from wild-type AAV2. The dashed line indicates equal ITR and EGFP titers. **(d)** QPCR titer for Set 1 variants. Scatter plot of mean log10 EGFP titer and mean log10 ITR titer for 100 Set 1 variants. Error bars show ±1 SEM of log10 titer across three technical replicates. Seven variants had fewer than 3 replicates for at least one titer. Mutation count was negatively correlated with both ITR titer (Spearman rho=-0.345, p=4.33e-04) and EGFP titer (Spearman rho=-0.483, p=3.65e-07). Log ratio of EGFP to ITR titer for payload-packaging variants is shown in **Figure S5a**. The log ratio of EGFP to ITR titer for low titer variants is not reported due to measurement noise at low titers. **(b)** QPCR titer for Set 2 variants. Scatter plot of mean log10 EGFP titer and mean log10 ITR titer for 155 Set 2 variants. Error bars show ±1 SEM of log10 titer across three technical replicates. Of 155 Set 2 variants, 68 are *virus-producing* (mean log10 ITR titer more than 3 SD above PBS), 23 of which are *packaging competent* (same threshold applied to log10 EGFP titer). Log ratio of EGFP to ITR titer for payload-packaging variants is shown in **Figure S5b**. (**f**) Transduction efficiency at 500 MOI in HEK293T cells for Set 2 variants. Bar chart of mean percentage of EGFP+ cells, with variants sorted and colored by mutation count against AAV2 VP3 (see Methods). Error bars are SEM across three biological replicates. 77 variants that had manufacture titer above 4.5x10^8 vg/ul are shown (one variant s2_53 was omitted). Wild-type AAV2 (dark green) and PBS (grey) are included as positive and negative controls. Bold x-axis labels mark the only variants (s2_129) with mean transduction efficiency above 50%. Two variants outside of these criteria were tested (**Figure S6b**).

### Design can alter cell type preference

VP3 surface residues can also determine tissue specificity. To evaluate whether generated sequences have altered specificity, transduction efficiency across cell lines from different tissues were used as an approximation. All variants from Set 1 included in the 500 MOI experiment (**Figure S6a**) and variants from Set 2 that transduced at least 1% of HEK293 cells were tested in HepG2 (liver), SH-SY5Y (brain), A549 (lung) and HEK293-Lenti-X (kidney) at 500 MOI (**Figure 4a**, additional variants excluded from the screening in **Figure S7**). Out of 19 variants of Set 1, 16 transduced HEK293 and SH-SY5Y cells with >5% efficiency and four variants achieved >50% transduction efficiency in both cell lines (**Figure 4b**). HepG2 and A549 were poorly transduced compared to WILD-TYPE AAV2 in all Set 1 variants, and altogether Set 1 showed homogenous cell specificity across variants.

Conversely, Set 2 variants showed multiple cell specificity profiles (**Figure 4c**). For instance, variants s2_132, and s2_0 showed >30% transduction of SHSY5Ys, but lower transduction in HEK293 cells (<10%). Variant s2_132 produced minimal transduction efficiency in every other line, while variant s2_0 robustly transduced HepG2s (>50%), A549s (>60%), and SH-SY5Ys (>40%), but not HEK293. Variant s2_129 showed very similar cell type preference to Set 1 variants, with non-significant differences between transduction in HEK293 and SH-SY5Y cells, but drastically reduced transduction efficiency in A549 and HepG2. Notably, variants s2_0 and s2_129 (both carrying >90 mutations) transduce SH-YSY5Y cells efficiently, but otherwise differ in tissue specificity, suggesting the model has encoded the capsid protein functional diversity seen in natural sequences. It was noted that most variants in Set 2 and all variants of Set 1 transduced HepG2 and A549 inefficiently. This could be caused by an Arg-to-Ser mutation in position 585 present in all Set 1 variants. However, only 21 of 155 Set 2 variants do not have a R585 mutation. Of these only seven were of sufficient titre to be included in the screening described in **Figure 3f** and none transduced >1% HEK293 cells. They were therefore not tested in the cell line experiment, meaning this hypothesis could not be tested with the current set.

**Figure 4.**
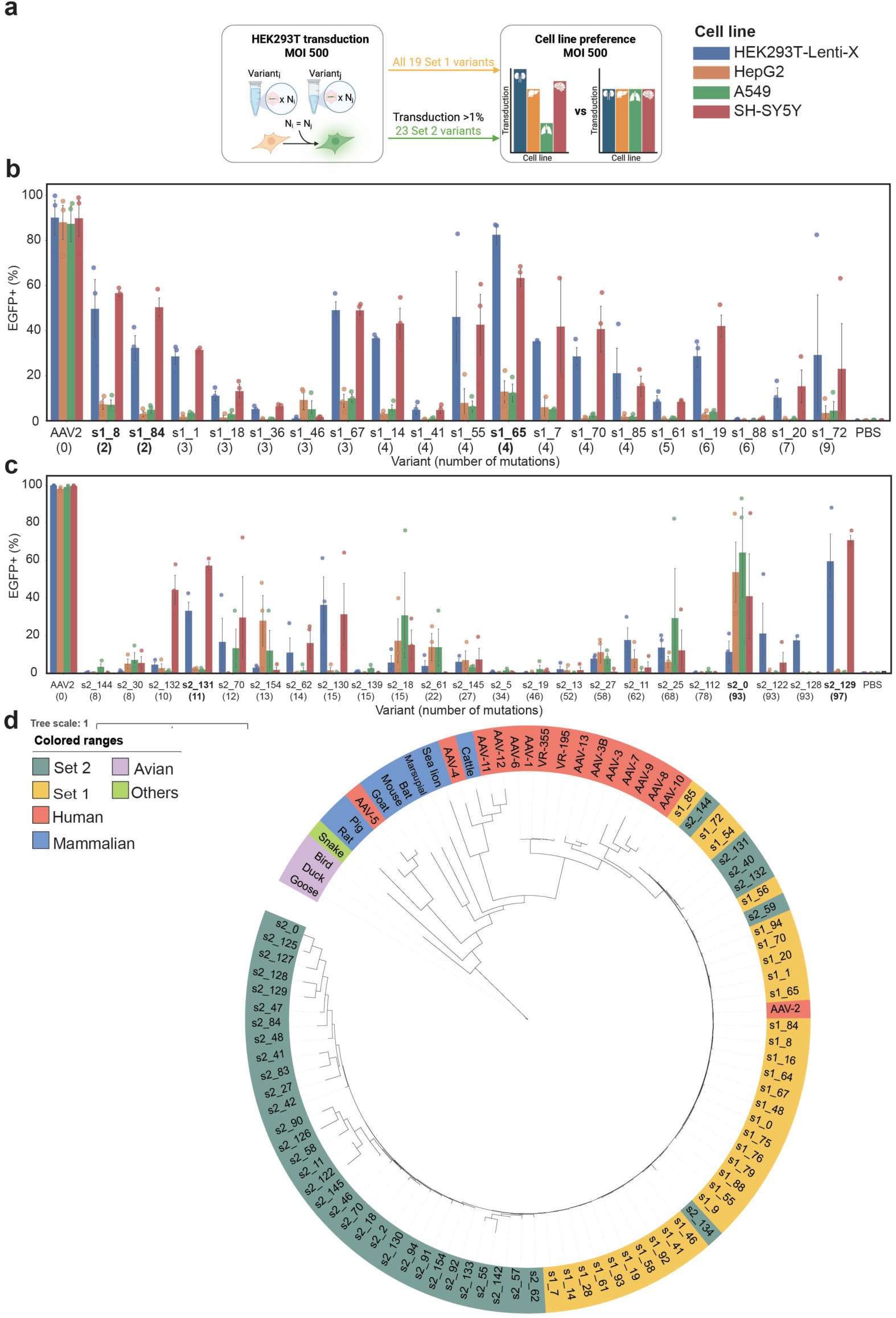
Validation of synthetic variants across multiple cell lines reveals two functional variants phylogenetically far from natural AAVs. (**a**) Selection criteria for cell line validation experiment. All variants of Set 1 tested at 500 MOI were also included in the cell line experiment, whereas only variants from Set 2 that transduced >1% HEK293 cells in the 500 MOI experiment (Figure 3f) were included (23 variants). (**b**) Transduction efficiency of Set 1 VP3 variants in multiple cell lines (MOI = 500). Each bar shows mean percentage of EGFP+ cells per cell line per variant, grouped by variant, colored by cell line. Error bar shows SEM across three biological replicates. Variant mutation count against AAV2 VP3 (see Methods) are shown in bracket under variant label. Wild-type AAV2 (first group) and PBS (last group) were used as positive and negative controls, respectively. Bold labels indicate the three variants with higher than 50% transduction efficiency in at least one cell line. **(c)** Multi-cell-line transduction efficiency of selected Set 2 capsid variants. Bar height indicates the mean %EGFP+ across replicates; error bars denote SEM. X-axis labels show variant identity and capsid mutation count. Bold labels indicate variants achieving greater than 50% EGFP+ in at least one cell line. Two recurring profiles emerged: preference for HEK293 (s1_65, s1_75, s2_128 and s2_130) or neuronal SH-SY5Y cells (s1_8, s1_84, s1_14, s1_19, s1_70, s2_131, s2_129 and s2_132); HepG2 liver and A549 lung cells were consistently least transduced. **(d)** Cladogram of representative natural VP3 sequences and functional Set 1 and Set 2 variants. The most highly mutated active variants form a tight cluster distinct from natural serotypes, indicating exploration beyond natural sequence diversity.

### Phylogenetic reconstruction suggests diversification beyond natural sequences

To understand where the generated functional sequences are within natural AAV sequence space, variants capable of transducing cells from both sets were used in a phylogenetic analysis with representative natural VP3s (**Figure 4d**). Set 2 variants formed a cluster with longer branches, both diverging substantially from natural AAV sequences and from each other. This cluster included variant s2_129, the highest-transducing set 2 variant, and variant s2_0, transducing ∼40% SH-SY5Y cells. Both differ from AAV2 at approximately 18% of VP3 residues. Notably, s2_0 and s2_129 differ from each other by 102 mutations, highlighting the capacity of generative models to reach distant functional sequences space beyond the neighborhood of natural sequences.

### Mapping mutations onto the capsid structure reveals evenly distributed surface change

Changing VP3 amino acid sequence by 18% is not by itself sufficient to broadly avoid recognition by nAbs with diverse epitopes. Even with a large number of mutations on exterior capsid surface, if they are clustered to one region, nAbs with epitopes in other regions might still neutralize the capsid. To evaluate the surface distribution, we reconstructed the full three-dimensional AAV2 capsid structure using PDB ID 7RWT and mapped designed AAV mutations onto it. **Figure 5** shows all 97 mutated amino acids from variant s2_129 displayed on the surface. Mutations are broadly distributed across the capsid exterior, likely altering most if not all antibody-accessible regions. This provides a plausible structural rationale for reduced recognition by nAbs with diverse epitopes broadly across different individuals.

**Figure 5.**
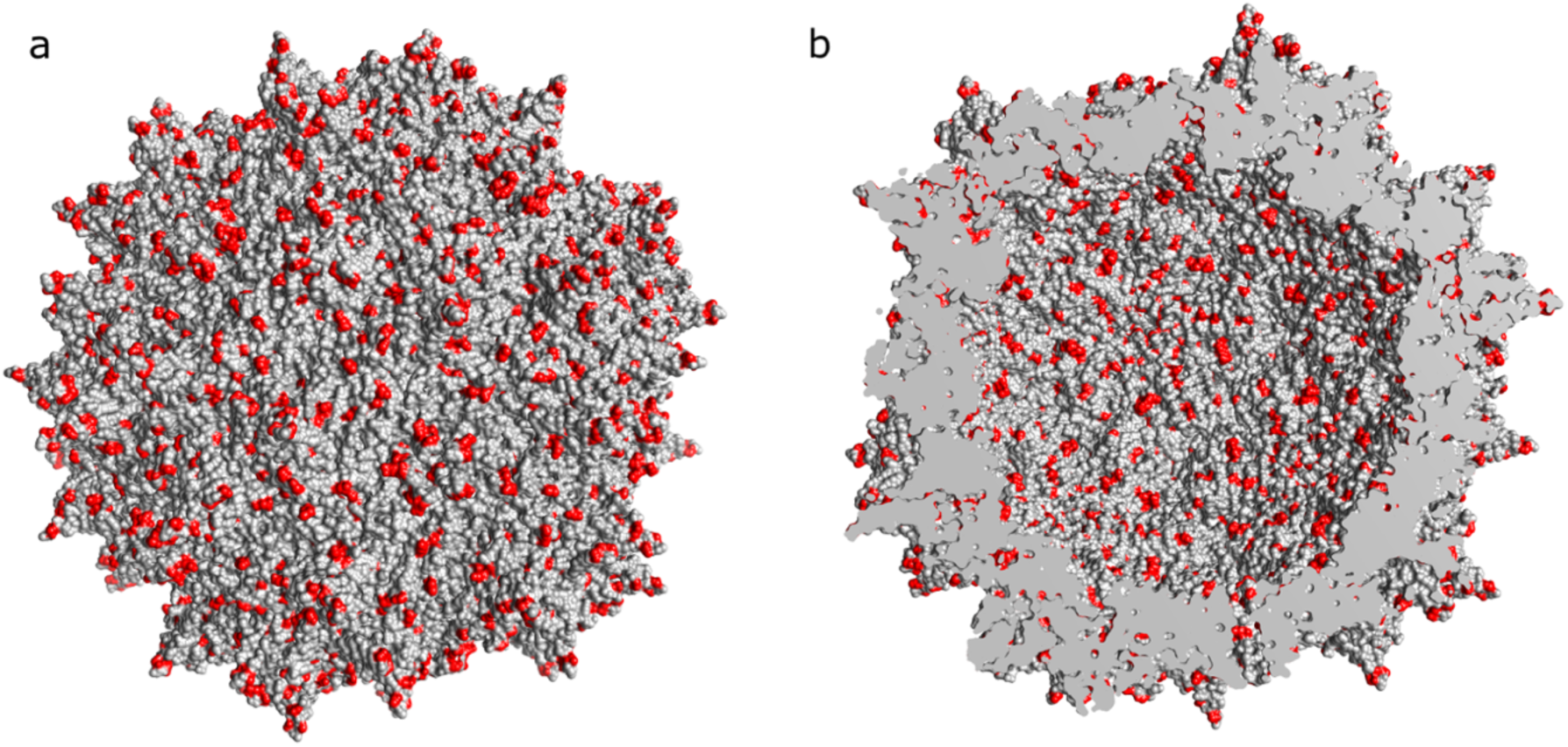
Distribution of designed mutations across the AAV2 capsid. The complete capsid was reconstructed from the AAV2 structure (PDB ID: 7RWT) and visualised using UCSF Chimera. The 97 VP3 residue positions mutated in variant s2_129 are highlighted in red across the capsid subunits; unmutated residues are shown in grey. Mutations are evenly distributed at both exterior (a) and interior (b) surface of the capsid.

### Synthetic AAV variants evade pre-existing neutralizing antibodies

To establish if synthetic variants avoid pre-existing immunity, human serum samples were screened for neutralization of AAV2 and 4 other clinically relevant serotypes (AAV2, AAV5, AAV8, AAV9 and AAVrh10). Serial dilutions from 20 different serum donors were tested for each serotype and NT50 values were calculated for each serum–serotype pair (**Figures 6b,c**). AAV2 was neutralized strongly by Donors 2 and 6, and moderately by Donor 16 (**Figure 6b**). Donors 2 and 16 also moderately neutralized AAV9 and AAVrh10; only donor 2 neutralized AAV5 and none neutralized AAV8 (**Figure 6c, S8a**). Variants from Set 1 and Set 2 were selected for neutralization evaluation according to their transduction efficiency (**Figure 3b,c**). For Set 1, variants transducing >30% cells were considered for neutralization experiments (27 variants). Of these, three variants (s1_09, s1_36, and s1_46) did not reach sufficient titre to be tested. Six randomly selected additional variants with <30% transduction efficiency were included as additional controls. For Set 2, the two highest transduction efficiency variants (in HEK293T) (MOI-matched, s2_131: 30.21%, s2_129: 64.18%) were selected, and another Set 2 variant was included as control (s2_92).

Sera from donors 2, 6 and 16 were then used to test all selected 33 variants (biological replicate n = 1, **Figure S8b**). Clinical exclusion thresholds range from an NT50 of 1:50 for intramuscular administration to <1:5 for systemic liver-directed delivery^26^. By this criterion, Donor 2 strongly neutralized wild-type AAV2, maintaining near-complete neutralization at low dilutions and yielding an NT50 of approximately 1:1,000. By contrast, all synthetic variants tested were neutralized less efficiently, with 28 variants yielding NT50 <1:50 in all three donors and 11 with NT50 <1:5 in all three donors. Notably, s1_72, s1_20, s1_41 and s2_129 resisted neutralization at every serum concentration tested. To validate some of the hits, we selected the two variants of Set 2 and randomly sampled 7 other Set 1 variants for further testing in three independent experiments per serum donor (biological replicate n = 3, **Figure 6d**). Representative curves compared wild-type AAV2 with variants s1_75 (5 mutations), s1_72 (9 mutations) and s2_129 (97 mutations) in Donor 2 serum (**Figure 6e**).

In summary, screening 255 variants across two sets identified four variants that transduce cells with >50% efficiency and also resist neutralization (NT50 < 1:50) of three different AAV2 positive donors (seven variants that transduced over 40% of cells). s2_129 was confirmed to resist neutralization in all three donors. S2_129 variant has ∼18% sequence diversity from the nearest natural sequence (AAV2). Its mutations are distributed across the entire VP3 region, including each of the Hyper Variable Regions (HVR), suggesting that thoroughly modifying exposed loops can facilitate evasion of neutralization by pre-existing nAb.

**Figure 6.**
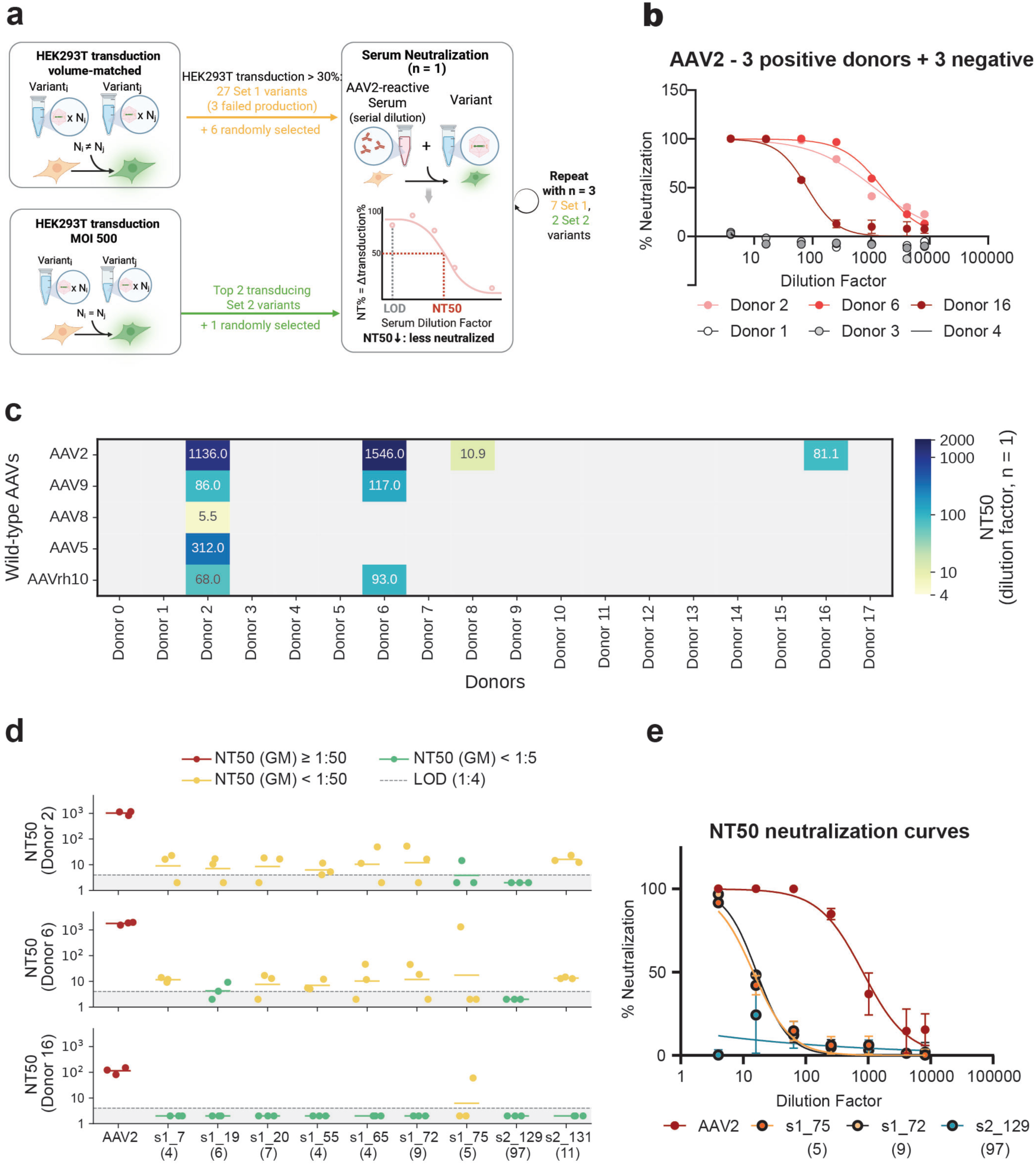
Synthetic variants resist neutralization by AAV2-positive sera. Neutralization was measured across serial serum dilutions, with dilution factor on the x-axis and NT50 calculated as described in **Methods**. Values are means ± SEM from three technical biological replica (*n* = 3) in technical triplicate per donor, unless specified. (**a**) Criteria of selection for neutralization assay. (**b**) Representative AAV2 curves for strongly positive donors 2 and 6, moderately positive donor 16 (red), and negative Donors 1, 3 and 4 (grey); all curves are shown in **Figure S8a**. Points show mean neutralization and solid lines the fitted curves. (**c**) NT50 neutralization experiment of five commercially available AAV serotypes (AAV2, AAV5, AAV8, AAV9, AAVrh10) against sera of 16 donors. Experiment in technical triplicate, biological replicas *n* = 3. (**d**) Bar plot representing mean NT50 values for variants selected from the initial screen (**Figure S8b**), ranging from low (white) to high (dark blue); each datapoint is the average of three technical replicas per single experiment. Points in red have NT50 > 1:50 and are considered neutralized, points in yellow have NT50 < 1:50 and could be considered for intramuscular administration. Last, teal points have NT50 < 1:5 and are considered to avoid entirely neutralization, being therefore eligible for systemic delivery in AAV2-positive patients. (e) Neutralization curves for wild-type AAV2 (dark red) and s1_72 and s1_75, and s2_129 (progressively darker purple with increasing mutation number) against Donor 2, which is positive for AAV2, but negative for each of the represented variants.

## Discussion

Generative protein models can propose sequences not sampled by nature that, with experimental selection, can yield variants with improved stability and function^27,28^. Here, ProteinVAE^18^ was adapted to model the AAV VP3 sequence and generate novel AAVs that avoid recognition by pre-existing immune responses. This required expanding the sequence exploration onto exposed surface residues and flexible loop, such as the HVRs. Earlier experimentally validated machine learning and generative AAV studies largely targeted short capsid segments only^13^, whereas recent proteinMPNN work re-designed primarily buried core and interface residues and kept nAb-facing surfaces unchanged^16^. Both do not address the problem of avoiding pre-existing antibody response.

The relatively small training set of natural AAV sequences introduces a challenge in learning the sequence space of its capsid proteins, because of the marked imbalance towards AAV2 sequences. This can potentially bias learned representation and reduce separation among close sequences from different serotypes. To mitigate it, contrastive representation learning was integrated to learn high-quality representation, capturing detailed differences between serotypes which are important for pre-existing antibody avoidance. These improvements yielded ProteinVAEc. With computational analysis, ProteinVAEc showed better recovery of general VP3 sequence and secondary structure patterns, and produced a smoother latent space in the important AAV2 region with improved sequence plausibility.

To validate ProteinVAEc experimentally and to generate useful variants, two variant sets were designed and tested. For validation, Set 1 sampled the near proximity of the latent space of AAV2, introducing up to 10 mutations. In a first screening, 19 variants transduced cells with efficiency higher than 50%, with 7 reaching efficiencies of >80%. When transduction was repeated at fixed viral titre (MOI = 500), only 4 variants transduced >50% cells. Set 2 was more exploratory, with variants containing 5 to 99 mutations. It yielded only one variant transducing >50% cells. 24 variants in total transduced HEK293T cells with a >1% efficiency. Transduction-competent sequences that were identified in Set 2 had up to ∼18% sequence diversity compared to the closest natural sequence.

When we investigated cell line preference among transduction competent cells, we noticed that all variants of Set 1 and most variants of Set 2 had lower specificity for HepG2 and A549 cells than wild-type. Variants in Set 2 transduced cell types with distinct preference profile. For instance, variants s2_154 and s2_61 disfavored HEK293 and SH-SY5Y for HepG2 and A549, s2_11 was selective for HEK293 cells, s2_132 showed higher selectivity to SH-SY5Y cells and s2_0 showed increased promiscuity. Overall, specificity for HepG2 and A549 cells was lower than AAV2 wild-type. This could be partially caused by the over-representation of mutation R585S, known to cause liver de-targeting^29–31^. All variants of Set 1 and 129 out of 155 variants of Set 2 carry a mutation at either R585 or R588^32^. These two arginine residues mediate the binding to heparan sulfate proteoglycan (HSPG) in AAV2 lab strains, which are a likely adaptation of cell culture propagation^33^, but are absent in circulating AAV2 and other AAVs isolated from human tissue^34,35^. ProteinVAEc training sequences are natural AAVs, which is why most of our designs do not have arginines at both positions. This also means in vitro experiments might underestimate the variants’ transduction for the more important in vivo setting.^27^

Neutralization experiments were possible in up to 33 variants using serum of three independent donors already positive for multiple natural AAV serotypes, including AAV2. 13 variants resisted neutralization across all three donors, with one variant (s2_129) completely resisting neutralization in the validation round. As might be expected, variants closer to AAV2 were more likely to be neutralized by sera positive for this serotype, while variants with higher number of mutations better resist neutralization. This was demonstrated in the validation round, where all variants selected for validation had NT50 < 1:50 (potentially eligible for intramuscular administration). As hoped, heavily modified capsids avoided neutralization by an even larger array of nAbs, illustrated by s2_129 which completely resisted neutralization across all sera and all dilutions tested (**Figure 6d, e**).

Interestingly, variant s2_129 also represents one of the two most diverse variants that successfully transduced cells. Indeed, variants s2_129 (97 mutations) and s2_0 (93 mutations) cluster at large phylogenetic distance from natural sequences and are themselves separated by 102 mutations. This highlights that generative models can reach distant functional sequence space when trained on limited number of natural sequences, a significant step toward addressing the redosing problemin gene therapy^36,37^

This data strongly reinforces the idea that applying generative models to synthetic AAV design can produce variants that avoid pre-existing neutralizing antibodies, thus expanding the reach of gene therapy to serve a large population of patients currently excluded due to their immunological profile. The discovery of multiple neutralization-resistant variants also offers the opportunity to explore their potential for sequential deployment and redosing, currently one of the most important and pressing unsolved challenges of the gene replacement therapy field.

### Limitation of the study

Despite the success in designing entirely AI-generated variants that effectively avoid neutralization, the screening and selection process highlighted a series of shortcoming of the novel variants. qPCR experiments showed a tendency of synthetic variants to carry partially packaged payloads, often causing low titres. This should be the next area of focus to advance towards clinical translation^38^. A potentially related issue is the reduction in transduction efficiency of most variants compared to AAV2 wild-type. Both issues likely arise from changes that disrupt interactions with the assembly machinery, such as the protein AAP that serves as a chaperon for nucleating assembly, or the packaging machinery composed of Rep proteins. It has been reported that partial packaging is reflected in lower transgene expression, without necessarily impacting transduction efficiency^25^. The training set is composed of sequences that may have introduced historical biases, for example variants carrying a Ser in position 584 of VP1 are frequent in natural AAV isolates, but rare in laboratory isolated AAVs^14,33^ . All of these limitations are ultimately a consequence of the lack of sequence data and annotations on transduction efficiency, cell type preference, and packaging efficiency. Large libraries designed specifically for efficient model training would be useful to advance this avenue of scientific investigation and innovation.

## Methods

### Computational Evaluation

To achieve a balance between diversity and quality, 1,500 vectors were sampled from a normal distribution with mean and variance of natural sample latent embeddings and decoded to new sequences. To maintain high sequence quality, sequences were ranked according to likelihood, and the top 1,000 sequences were kept for analysis. ProGen2-small (151M) was fine-tuned on the same VP3 dataset as a comparison. 1,500 sequences were generated with prompt “M”, and the top 1,000 sequences with highest likelihood were kept.

To analyze sequence-level features, generated sequences and natural sequences were aligned in multiple sequence alignment (MSA). After removing columns with low occupancy (80%), per-column Jensen–Shannon divergence (JSD) was calculated between amino acid distributions in generated sequences and natural sequences. Lower JSD indicates closer agreement with natural evolutionary constraints at each position.

For assessing structural features, 3-state secondary structure was predicted from sequence using SPOT-1D-Single^43^. Helix and strand fractions were compared between natural and generated sequences to evaluate whether generated sequences reproduced the secondary structure composition of natural VP3 proteins.

Latent space interpolation was performed between AAV2 and AAV1, two serotypes that can form mosaic capsids^19^. Hamming distance to each parental sequence was computed along the interpolation path to assess whether the latent space supports smooth traversal between compatible serotypes. The Hamming distance is calculated by padding both sequences to same length, then dividing number of different positions by length. Likelihood of interpolated sequences was evaluated using the fine-tuned ProGen2 model as an independent evaluator.

### Phylogenetic Analysis

Amino acid sequences of the viable Set1 and Set2 AAV VP3 were aligned to representative natural sequence using Clustal Omega with default parameters. Phylogenetic reconstruction was performed using the Maximum Likelihood (ML) method implemented in PhyML version 3.^44^. Briefly, the LG amino acid substitution model was utilized, with equilibrium frequencies estimated by the model. Rate heterogeneity across sites was accommodated using a discrete Gamma distribution with four rate categories (+G) and an estimated shape parameter (*α*), alongside an estimated proportion of invariable sites (+I). Initial tree construction was performed using the BioNJ algorithm. To ensure rigorous exploration of tree space and avoid local optima, tree topology and branch lengths were optimized simultaneously using the Nearest Neighbor Interchange (NNI) heuristic search algorithm supplemented with five random starting trees. Statistical confidence of the branch nodes was evaluated using standard bootstrap analysis with 500 replicates. The final consensus tree was visualized and annotated using iTOL^45^.

### Sequence Generation and Selection for Experimental Validation Set 1: Near-AAV2 Variant Generation and Filtering

A total of 4.5 million latent vectors were sampled from a Gaussian distribution centered at the AAV2 latent representation with standard deviation of 2. The top 3 million vectors by model likelihood were retained. In parallel, 1,000 latent vectors were generated by linear interpolation between AAV2 and serotypes 1, 3, 7, 8, and 9, which have been reported as compatible with AAV2^19,32,46^. After decoding, 2.7 million unique sequences were obtained from free generation and 689 unique sequences from interpolation.

A series of filters was applied to enrich for functional candidates. First, sequences carrying double mutations at R585 and R588 were excluded because this combination, while potentially altering tropism, has been reported to abolish transduction in HEK293 cells^47^. Next, an activity filter was applied using previously published single-mutation mouse biodistribution data: for each candidate sequence, the sequence-level selection score was computed as the mean of the selection scores of all its single mutations^14^. Any sequence showing less than a 1.8-fold increase relative to wild-type across all organs was removed, as it is less likely to be functional. As the effect of mutation is less likely to be additive at higher number of mutations, only sequences with fewer than 10 mutations were kept. In addition, candidates with mutations at residues involved in AAV2 AAVR interaction were excluded^48^. Finally, to maintain diversity in the selected sequences, leader selection ^49^ was used to select the final set of 100 variants, maximizing coverage of distinct mutation positions.

### Set 2: Uncertainty-Guided Exploration of Distant Sequence Space

To explore unsampled regions of AAV VP3 sequence space, 150,000 latent vectors were sampled from a Gaussian distribution centered at the AAV2 latent representation with an increased standard deviation of 3. The 50,000 vectors with the highest model likelihood were retained and decoded. Sequences carrying 5–99 mutations relative to AAV2 were selected for downstream prioritization.

#### Uncertainty-based prioritization

To identify candidates expected to be most informative for model refinement, predictive uncertainty was estimated using Monte Carlo dropout. The 3,000 sequences with the highest uncertainty were retained as the initial candidate pool.

#### Activity classifier

Because mutational effects are unlikely to remain additive beyond approximately ten substitutions, a viability predictor was established by fine-tuning ESM2 on natural VP3 sequences and single-mutation sequences^49^ The 3,000 uncertainty-selected sequences were evaluated using this predictor. Briefly, the 650M-parameter ESM2 model was adapted for binary sequence classification (viable versus non-viable) on full-length natural VP3 sequences (augmented with 5-20 random mutations as negative samples), our Set 1 packaging results, and previously published packaging data of AAV2 single-mutation sequences ^14^. Parameter-efficient fine-tuning was performed with low-rank adaptation (LoRA; rank 4, *α* = 1) applied to the query, key, value, and dense projection matrices, while the classification head and all bias terms were left trainable. The model was trained for 20 epochs with early stopping on validation loss. AdamW optimizer with a learning rate of 3 × 10^−4^, an effective batch size of 25, and a maximum sequence length of 1024 tokens. On the held-out validation set the predictor reached a precision of 0.73.

#### Diversity- and activity-aware selection

To ensure balanced representation across diversity and viability, multiple complementary selection criteria were applied. First, 10 sequences with the highest uncertainty were retained to maximize potential information gain. Next, the 10 top variants by predicted viability were included to ensure representation of highly functional candidates. To encourage exploration across mutation ranges, sequences were stratified by total mutation count relative to the closest natural VP3 sequence, and 10 high-viability candidates were selected from each of the 10 bins. Finally, to maintain a pool of moderately mutated but viable variants, sequences with 5–30 substitutions were filtered, and the top 75 sequences, ranked by predicted activity, were retained.

In total, 155 variants were selected for Set 2, with some overlapping variant between the 3 prioritization methods described above. This staged prioritization process yielded a set of candidates spanning a broad mutational range while enriching for both informative and predicted high-viability sequences, supporting balanced exploration of AAV VP3 sequence space. However, the overall exploration success rate relies heavily on the activity predictor (see Discussion).

### Cloning

An Acceptor Plasmid was designed and built based on pCap2Rep2 plasmid (Takara), where the region corresponding to VP3 (residues 203-735 of VP1) of AAV2 was replaced by a GFP drop off cassette, flanked by inverted PaqCI sites for negative selection. All designed variants of VP3 coding region were synthesized by GenScript™ (Piscataway, NJ, USA), without further codon optimization. Designed variants of Set 1 and 3 were received as inserts in a pUC57 plasmid and then subcloned into the Acceptor Plasmid by PCR amplification (Q5 polymerase, New England Biolabs, Ipswich, MA, USA, E1601) and subsequent Golden Gate cloning using PaqCI (New England Biolabs, Ipswich, MA, USA, E1601). During the PCR amplification, primers were designed to include complementary PaqCI sites to enable scarless Golden Gate cloning.

Variants of Set 2 were subcloned by GenScript™ (Piscataway, NJ, USA) in correspondence of region 203-735 of VP1 in the coding region for the Cap gene of a proprietary RepCap plasmid.

Variants of Set 4 were subcloned by Golden Gate, by designing 5 fragments each flanked by a different PaqCI site for scarless combinatorial assembly. The sequences of wild-type AAV2 VP3 and that of s2_129 were divided in 5 regions, respectively denominated N-terminal (N), packaging 1 (P1), Central (M), packaging 2 (P2), and C-terminal (C). The crossover point between adjacent regions was designed to retain sequence identity across regions in a span of 6bp, necessary for designing Golden Gate overhangs. Each fragment from AAV2 wild-type and s2_129 was PCR-amplified with primers including PaqCI overhangs defined for each specific position, or custom synthesized (GenScript™). PaqCI sites were designed to reduce spurious recombination^50^. Ultimately, fragments were combined with the Acceptor Plasmid in a 1:1:1:1:1 ratio for Golden Gate reaction (New England Biolabs, Ipswich, MA, USA, E1601).

All plasmids were transformed and propagated in NEBStable *E.coli* cells (New England Biolabs, Ipswich, MA, USA, E1601) at 30^◦^C.

### Cell Culture

HEK293T Lenti-X cells were propagated in Dulbecco’s Modified Eagle Medium (DMEM, Gibco, 11995073) 10% Fetal Bovine Serum (FBS, Gibco, A5256701) media, with 1% Penicillin Streptomycin Solution (Pen/Strep, Life technologies). Cell lines A549, HepG2 and SH-SY5Y were cultured in DMEM+ 10% FBS.

Passaging was performed by lifting cells with Trypsin (Gibco, 25200072) and resuspending them in a 1:5 or 1:10 dilution. Cell counting performed with a Countess automatic cell counter (Thermo Fisher). Lenti-X Cells with a passage number no higher than P=16 were used for viral production.

### Viral production (Set 1)

AAV particles were produced by triple transfection using adenoviral helper plasmid (pHelper, Takara), payload plasmid with self-complementary ITRs, expressing EGFP controlled by CAG promoter (scAAV-CAG-GFP, Takara), and AAV2 Rep/Cap (Takara) in case of wild-type positive control, or otherwise the plasmid of interest. Transfection was performed using Polyethyleneimine (PEI) as previously described^51^ in 6 well plates, with total media of 2ml/well. After 24 hours, transfection efficiency was validated by GFP expression and media was changed for Production Media as described in^51^. Harvest was carried out 5 days after media change, by adapting a previously published protocol^51^, which included a lysis step with citric acid. Supernatant was retained and combined with the viral solution after the extraction step. An overnight precipitation with polyethylene glycol (PEG) 8000 was adopted to concentrate the viral solution. Pellets were resuspended in 50 *µ*l of PBS +0.001% Pluronic™ F-68 buffer.

### Viral production (Set 2)

AAV particles were produced by GenScript™ (Piscataway, NJ, USA), by means of transient transfection using PEI. Briefly, 4 x 15cm dishes with Lenti-X 293T cells seeded at 1.8 × 10^7^ cells per dish, followed by cell lysis, PEG precipitation, and resuspension in 1ml of DPBS + 0.001% Pluronic™ F-68 buffer. Variants were quantified with qPCR by the supplier.

### Viral production (Neutralization assays, 500 MOI transduction)

AAV particles were produced by triple transfection using pHelper, scAAVCAG-EGFP and plasmid of interest (or pRepCap) in a T-75 plate (Sarstedt, 83.3911.002) as previously described^51^. One day after transfection, efficiency was assessed by GFP expression and media was changed to production media ^51^ at a constant volume of 10ml. After 5 days from media change, the supernatant was separated from cells. Cells were lifted with a cell scraper (Fisherbrand) and lysed in a citric buffer ^51^. After lysis and separation of the viral supernatant from the cell pellet, viral supernatant and exhaust cell media containing AAV were combined and filtered using a 0.45 *µ*m syringe-driven filter (13100107, Fisher Scientific). Clarified and filtered solution was then precipitated overnight with PEG8000 and resuspended in 100*µ*l 0.001% Pluronic F-68 PBS solution.

### DNAseI reaction

Prior qPCR or ddPCR quantification, samples were diluted 1:100 and an aliquot of these dilutions was mixed with a DNAseI reaction mix (Worthington, LS006355) according to provider recommendation. Reaction was carried out for 4 hours at 37^◦^C followed by heat denaturation at 95^◦^C for 10 min to release encapsulated AAV genomes.

### ddPCR

Samples were titrated through ddPCR targeting the EGFP transgene by The Center for Applied Genomics (TCAG, SickKids, Toronto, ON), or using a custom ITR probe obtained from ^23^.

### qPCR

Quantitative PCR was carried out in 10*µ*l reactions in triplicate with a Luna qPCR kit (New England Biolabs, Ipswich, MA, USA, E1601). In each reaction, either EGFP primer set (EGFP fwd: GGACGACGGCAACTACAAGA; EGFP rev: AAGTCGATGCCCTTCAGCTC), or the ITR primer set (ITR fwd: GGAACCCCTAGTGATGGAGTT; ITR rev: CGGCCTCAGTGAGCGA) was used ^23^.

qPCR for EGFP was carried out with a 1:1 ratio of forward : reverse primer, whereas qPCR for ITR was carried out with a forward : reverse primer ratio of 1:3.4, based on published protocols^23^. In either case, reaction was carried out in a Quant Studio 5 thermocycler (Thermo Fisher Scientific), in a 384 well plate Bio-Rad, HSP3801, sealed with transparent sealing film (FroggaBio, ATL-QPCR-100). Reaction conditions were 2 minutes at 50^◦^C, 10 minutes at 95^◦^C, then 40 cycles at, respectively 95^◦^C for 15 seconds and 60^◦^C for 1 minute. In each plate, a standard curve of 7 serial dilutions 1:5 from an independently measured wild-type AAV2 stock was prepared in triplicate for each primer set. Amplification curves were analyzed with the Standard Curve function on the Thermo Fisher Scientific Cloud platform to obtain Cq values. Conversion of Cq values into concentrations expressed as vg/µl was carried out by establishing a standard curve per plate, per each primer set and then interpolating each dCq value to the relative standard curve within the same plate (**Figure S6**). Downstream analysis and statistical analysis were carried out on Microsoft Excel and GraphPad Prism.

### Transduction experiments (screening and validation)

In both Set 1 and Set 2, HEK293T cells were seeded at 10,000 cells per well in 96 well flat bottom TC-treated plates (VWR, 10062-900). The next day, cells were transduced with 10*µ*l viral preparation, either freshly produced and resuspended (Set 1) or 10*µ*l of a dilution 1:10 from the original stock vial (Set 2), except for vial from variant s2_116 which was not diluted, due to low titer (10^6^ vg/ml). Given the second lowest titer among viruses in Set 2 was in the 10^7^ vg/ml order of magnitude (**Table S1**), this dilution granted at least MOI = 1 per variant. Transduction was carried out in technical triplicate, in three independent experiments (n=3 biological replicas). Statistical tests were carried out on logit transformed data (**Figure S3**): one-sided Welch’s t-test on log EGFP+%, BH q ≥ 0.05.

In the validation experiment with multiple immortalized cell lines, cells were transduced at a constant MOI = 500. In that case, viruses were diluted in 0.5 × 10^6^ vg/*µ*l in PBS+0.01% F-68 Pluronic in a 130*µ*l stock solution and then delivered as 10*µ*l/well in triplicates in each cell line. Cell lines used in the immortalized lines validation experiment were SH-SY5Y, A549, HepG2 and HEK293T. Each line was seeded at 10,000 cells per well and cultured in DMEM+10% FBS overnight prior transduction.

Infection was carried out for 72 hours after which cells were harvested in 96 well conical bottom plates (Sarstedt, 82.1583.001) and stained with live/dead™ Fixable Near-IR Dead Cell Stain kit (Thermo Fisher, L34975) for 5 minutes at room temperature. Reaction was quenched by diluting the staining solution 1:5 and rapidly pelleting cells. After staining, cells were fixed with 4% Paraformaldehyde (PFA) for 15 minutes at room temperature and conserved at 4^◦^C in the dark until measurement (1-36 hours later).

For Set 2, validation transduction at matching MOI (MOI=500) was carried out by calculating dilutions using supplier-reported titres (**Table S1**). Transductions were carried out in technical triplicate across three independent experiments, where a different vial was used each time.

### Flow cytometry

Flow Cytometry data recorded on a BD LSRFortessa and a BD High Throughput Sampler (HTS), with a FACSDiva Software (v6.2). FITC data (GFP) was recorded with a blue laser (488nm) and a filter set 530/30, 505LP; APC-Cy7 (Live/Dead staining) was recorded with a red laser (640nm) with a filter set 780/60 750LP. Data analyzed with FlowJo (v10.10.1) and GraphPad Prism (v10.2.3). Transduction efficiency was calculated as GFP%,

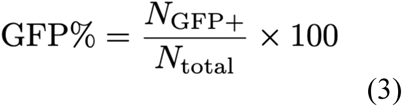

where *N_GFP_*_+_ is number of cells showing GFP in flow cytometry, *N_total_* is total number of cells sorted. EGFP+% was log-transformed as replicate standard deviation scaled approximately linearly with the mean (slope ≈ 1 on log–log plot; Figure S3), consistent with multiplicative noise). Each variant was compared with control by a one-sided Welch’s t-test on log-transformed EGFP+%. P-values across variants were adjusted by the Benjamini–Hochberg procedure, variants with adjusted q < 0.05 were considered significant. Variants with EGFP+% over 80% and not significantly lower than AAV2 were called high-transducing. Variant with EGFP+% significantly higher than PBS were considered transducing.

### ^54^Neutralization assays

HEK293T cells cultured in DMEM+10%FBS were seeded at a density of 10,000 cells per well and incubated overnight prior to transduction. Serum samples (obtained from StemCell Technologies, Cat#200-0160) were diluted with DMEM at the following concentrations: 1:4, 1:16, 1:64, 1:256, 1:1024, 1:4096, 1:8192. Transduction was carried out at final MOI = 500. To do so, virus samples were diluted to 1 × 10^6^ vg/*µ*l in PBS+0.01% pluronic acid. The diluted serum was mixed in equal volume with diluted virus in a round bottom 96-well plate and incubated at 37^◦^C for 1h. HEK293T cells were transduced with 10 *µ*l of serum : virus mixture in triplicate and incubated for 72h at 37^◦^C. After 72h, the cells were lifted and stained with Live/Dead-Near IR (Thermo Fisher, L34976) and fixed with 4% PFA.

Percentage of viruses neutralized by the serum (Neutralization% or NT%) is calculated as,

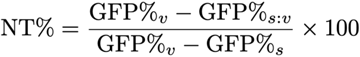

where GFP% denotes the transduction percentage (**Equation 3**). GFP%_v_ and GFP%_s_ are obtained with pure virus sample or pure serum, respectively. GFP%_s:v_ represents each serum-virus mixture. With this definition, NT% is 0% in pure virus (no serum), and 100% in pure serum sample (no virus).

Data was analyzed with GraphPad Prism (v10.2.3). For each donor and virus, a neutralization curve was obtained and NT50 was calculated from the nonlinear regression of log(inhibitor) versus response (variable slope). An NT50 value corresponds to the dilution factor where 50% of the virus is neutralized. Based on this rationale, variants positive for neutralization were defined by applying two cutoffs of, respectively NT50=50 and NT50=5, based on ^26,52,53^.

## SUPPLEMENTARY FIGURES

**Fig. S1.**
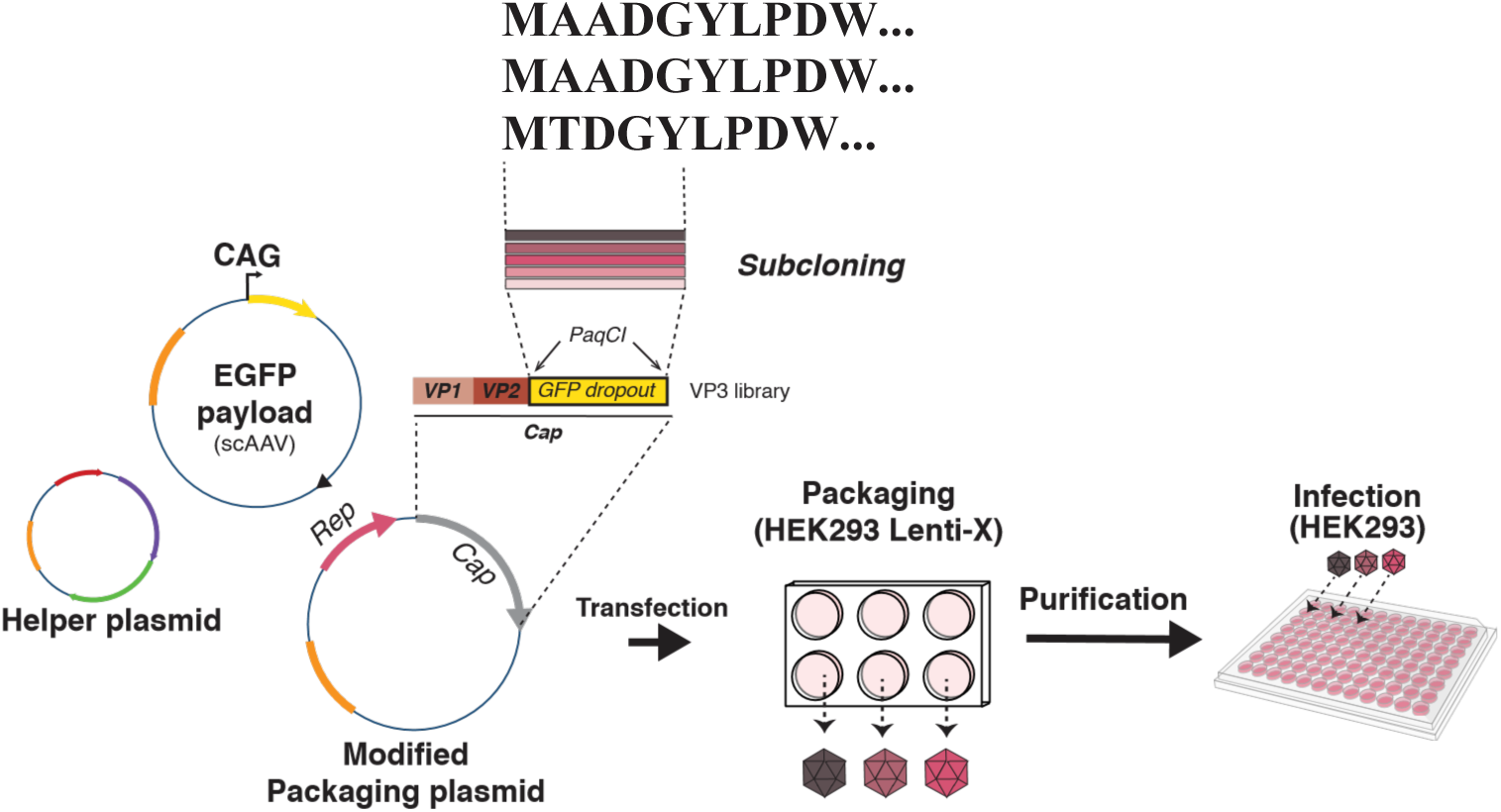
Plasmid map and construct design. The modified packaging plasmid was modified to host a PaqCI Golden Gate acceptor. A Golden Gate reaction replaces the GFP dropout cassette with a synthetic VP3. Cloned RepCap vectors are transfected along with a plasmid expressing the Helper genes and a payload plasmid, encoding a reporter EGFP driven by CAG promoter, flanked by self-complementing ITRs. Variants are purified and transduced separately.

**Fig. S2.**
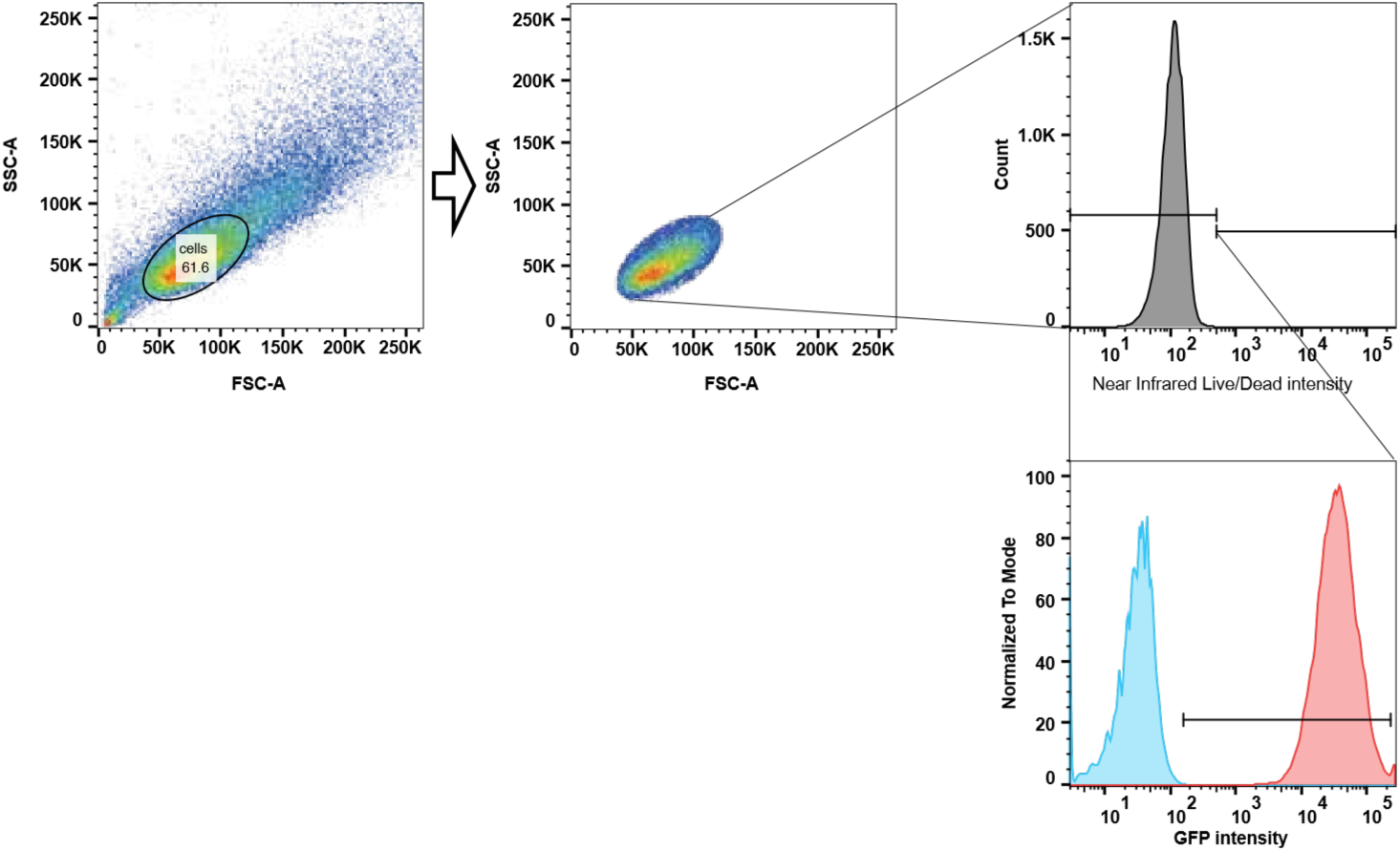
Gating strategy. A first gate was established to exclude cell aggregates and doublets. Then, single cells were gated according to Live/Dead staining, selecting the non-stained population (i.e. live population). The live cell population was then gated for GFP+, by comparing GFP fluorescence of non-transduced cells (NT, gray) with wild type (WT, red) cells.

**Figure S3.**
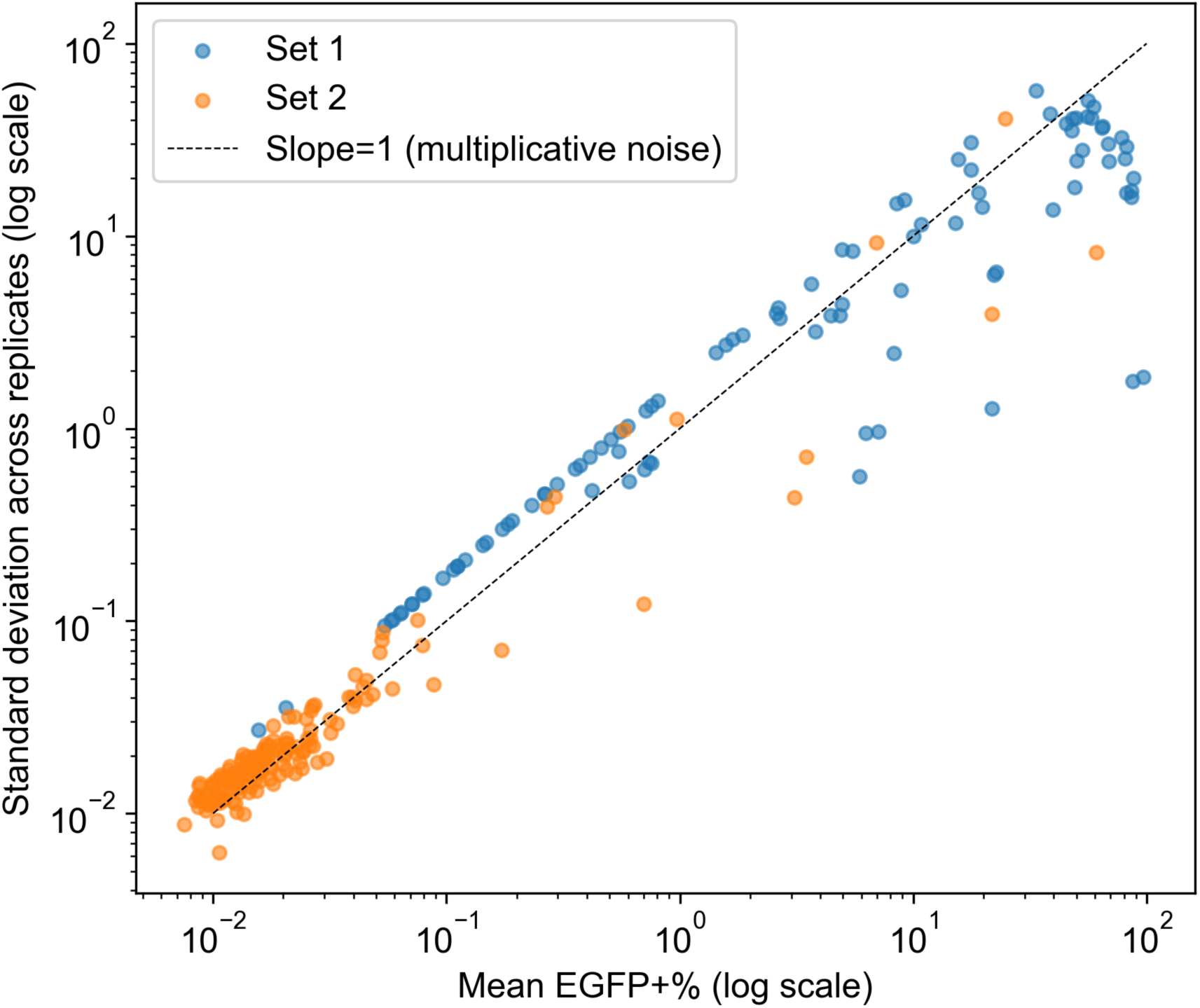
Replicate variation scales with transduction efficiency. Standard deviation across 3 replicates plotted against mean EGFP+% for every variant in both sets (log–log scale). The dashed line indicates slope = 1, the relationship expected under purely multiplicative noise. Variation across replicates increases as transduction efficiency increases, therefore, log transformation was applied before statistical testing (Methods).

**Figure S4.**
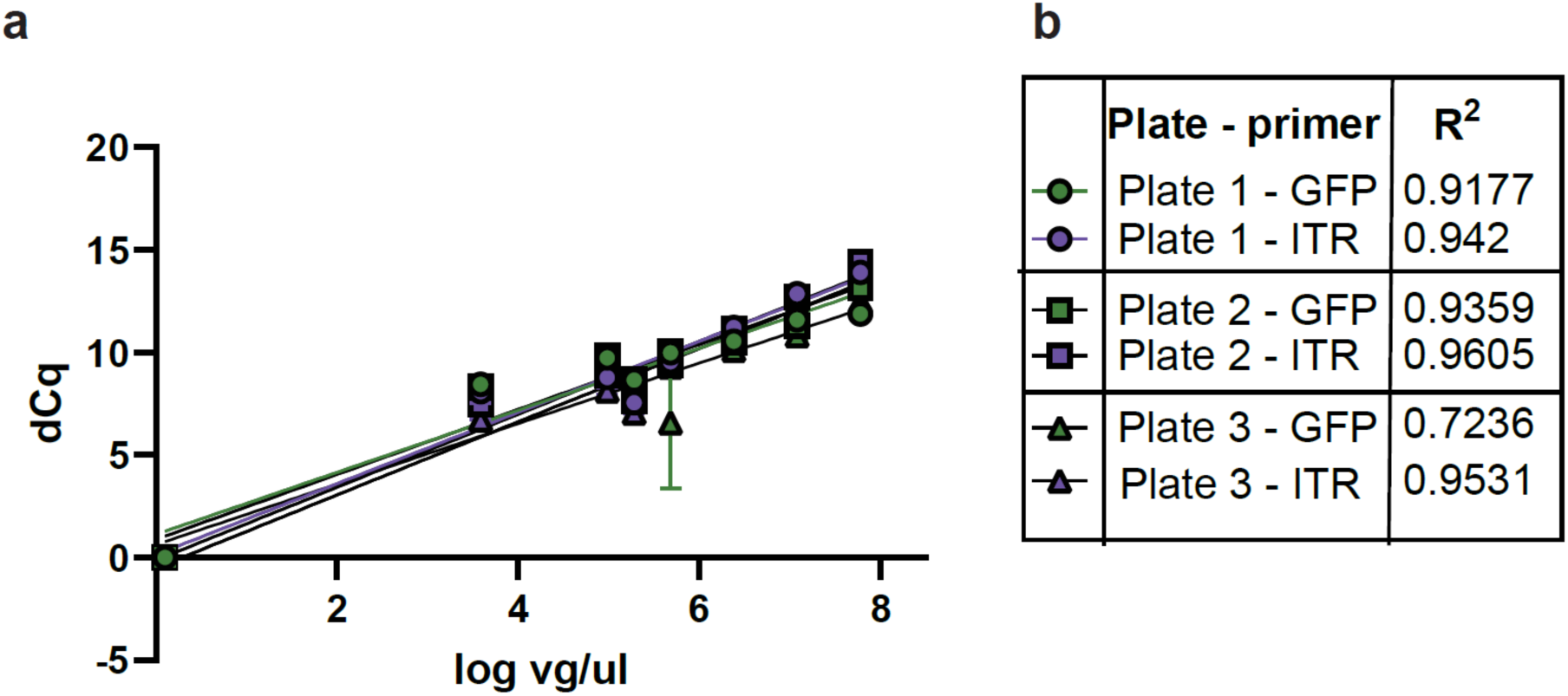
Representative standard curve used in the conversion of dCq values from qPCR into viral titers. (a) Serial dilutions 1:5 of a stock solution of commercial AAV2 were digested with DNAseI in parallel to the samples measured and quantified with both primer sets (shades of green for EGFP, shades of purple for ITRs). Values of dCq on the y axis are then reported to a concentration independently calculated on the x axis. Concentration of the stock solutions are based on ddPCR quantification and are expressed here in log_10_ of viral genome per microliter (vg/ml). A linear fit is then used to obtain a calibration curve. Measurements were taken in triplicate, and a separate set of standard curves was performed in each plate (darker shade of color as plate number progresses). Concentration of samples belonging to each plate were converted using the relative standard curve. (b) *R*^2^ values of the relative standard curves.

**Figure S5.**
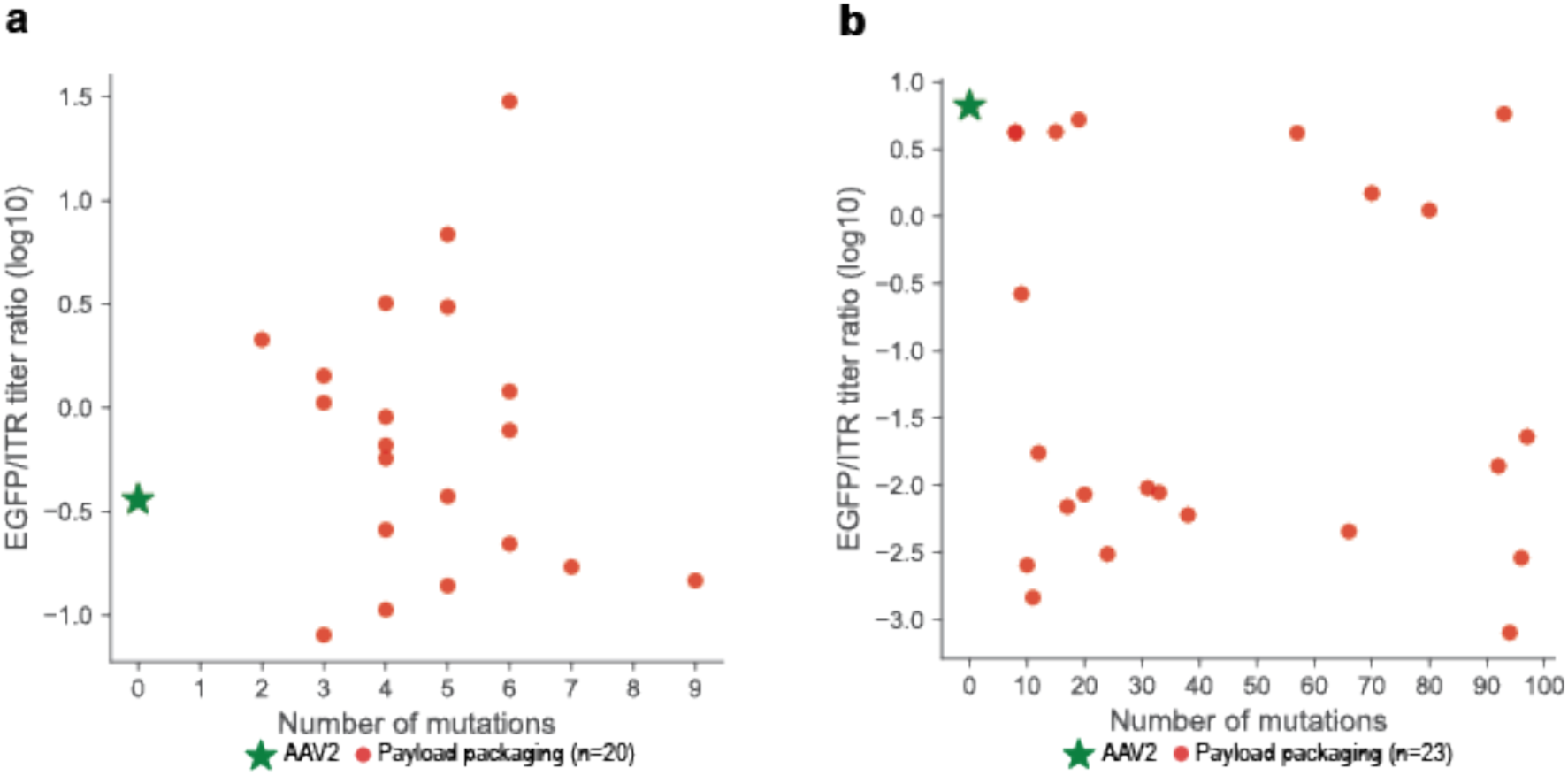
EGFP/ITR ratio per variant, ranked by number of mutations. **(a)** Log 10 EGFP to ITR titer ratio vs. mutation count for Set 1 payload-packaging variants. Scatter plot of mean log10 ratio EGFP/ITR titer against number of mutations from wild-type AAV2 VP3 (see Methods) for the 20 of 100 Set 1 variants classified as payload-packaging (mean log10 EGFP and ITR titers both more than 3 SD above PBS; red circle). Wild-type AAV2 (green star) is shown as a positive control. **(b)** Log 10 EGFP to ITR titer ratio vs. mutation count for Set 2 payload-packaging variants. Scatter plot of mean log10 ratio EGFP/ITR titer against number of mutations from wild-type AAV2 VP3 (see Methods) for the 23 of 155 Set 2 variants classified as payload-packaging (mean log10 EGFP and ITR titers both more than 3 SD above PBS; red circle). Wild-type AAV2 (green star) is shown as a positive control.

**Figure S6.**
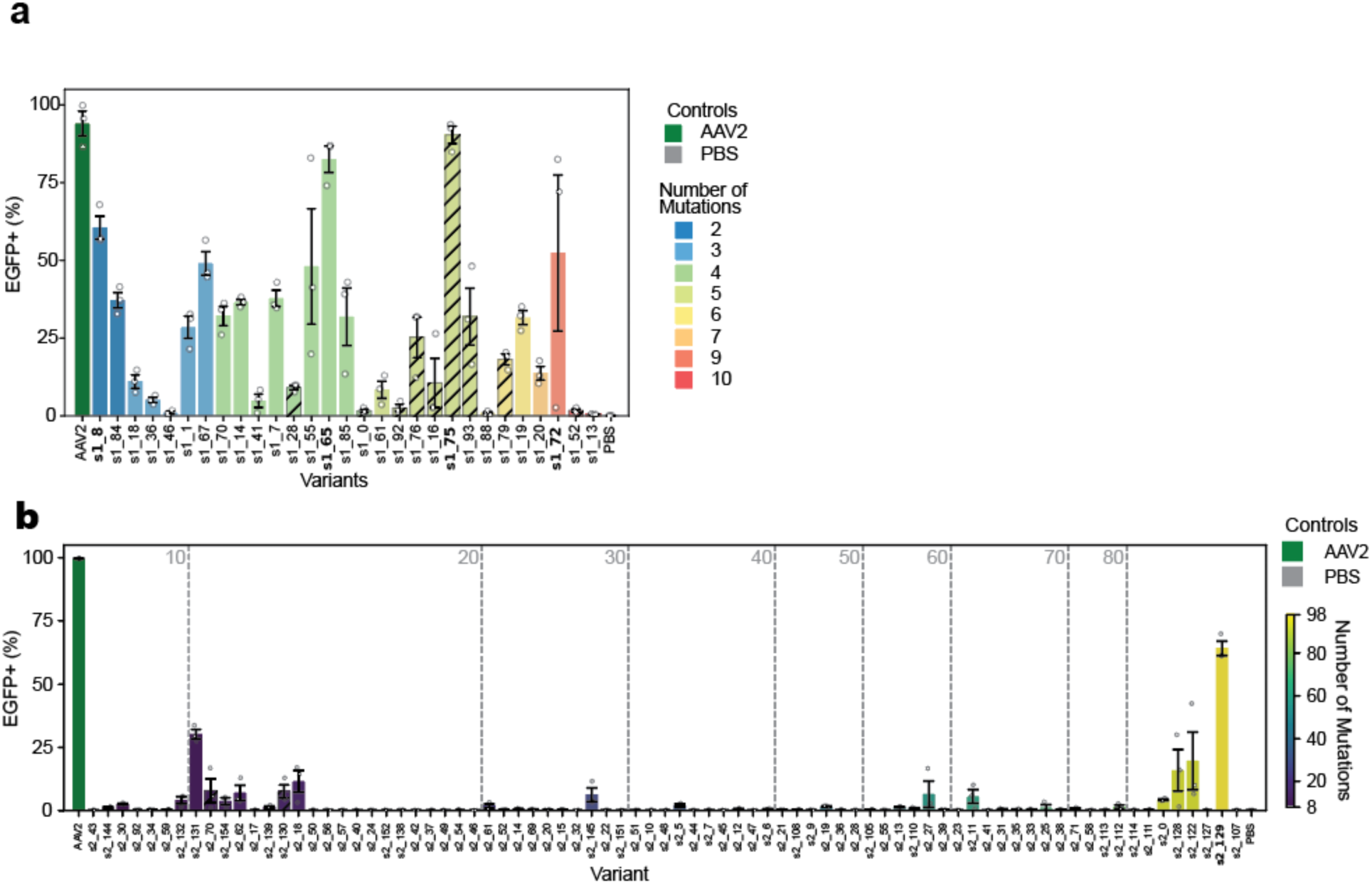
Extended data for transduction experiments at 500 MOI. **(a)** Transduction efficiency for Set 1 variants at 500 MOI Bar chart of mean percentage of EGFP+ cells, with variants sorted and colored by mutation count against AAV2 VP3 (see Methods). Error bars are SEM across three biological replicates. Hatched bars represent variants that did not transduce >50% in the original transduction experiment (**Figure 3b**) but had high titres, picked at random. (**b**) Extended data for transduction efficiency at 500 MOI in HEK293T cells for Set 2 VP3 variants. Bar chart of mean percentage of EGFP+ cells, with variants sorted and colored by mutation count against AAV2 VP3 (see Methods). Error bars are SEM across three biological replicates. 77 variants that had manufacture titer above 4x10^8 vg/ul are shown (one variant s2_53 was omitted). Wild-type AAV2 (dark green) and PBS (grey) are included as positive and negative controls. Bold x-axis labels mark the only variants (s2_129) with mean transduction efficiency above 50%. Two variants outside of these criteria were tested (s2_70, s2_130, hatched bar).

**Figure S7.**
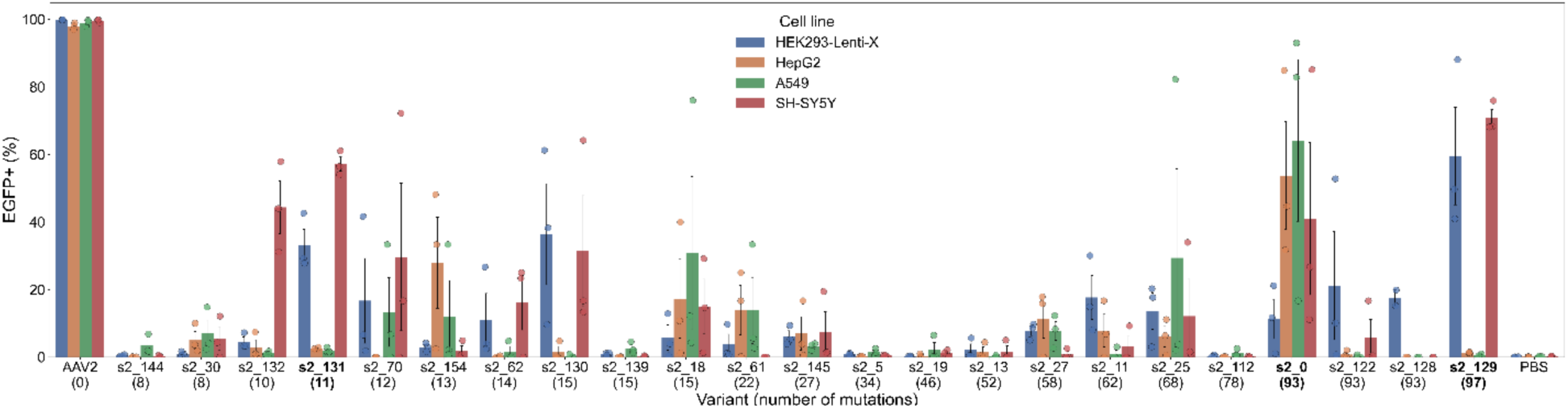
Extended data for transduction experiment in HEK293, HepG2, A549 and SH-SY5Y lines, Set 2. Extended data for transduction efficiency of Set 1 VP3 variants in multiple cell lines (MOI = 500). Each bar shows mean percentage of EGFP+ cells per cell line per variant, grouped by variant, colored by cell line. Error bar shows SEM across three biological replicates. Variant mutation count against AAV2 VP3 (see Methods) are shown in bracket under variant label. Twelve variants exceeding 50% transduction in the initial volume-matched transduction screening assay are shown in **Figure 2e**. Wild-type AAV2 (first group) and PBS (last group) were used as positive and negative control, respectively. Hatched bars show five variants that did not meet the screening criteria but were tested. Bold labels indicate the four variants with higher than 50% transduction efficiency in at least one cell line.

**Fig. S8.**
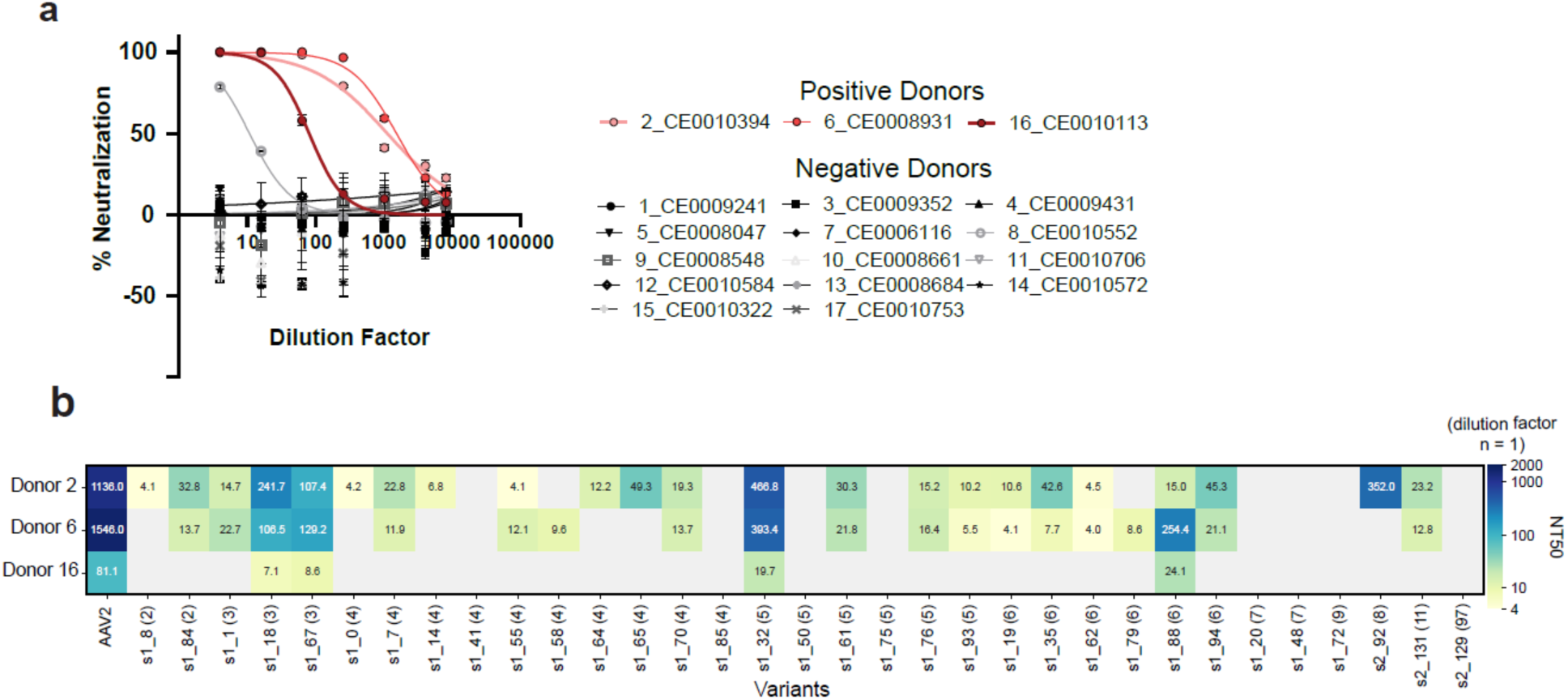
Most synthetic variants resist neutralization by serum of three separate donors, positive for AAV2. Neutralization assays represented in the form of neutralization curves and NT50 bar plots. **(a)** Neutralization curve of AAV2 against all sera tested. Experiment in technical triplicate, biological replicas *n* = 3. (**b**) NT50 neutralization experiment of 33 selected variants against sera of 16 donors. Experiment in technical triplicate, biological replica *n* = 1.

